# Endogenous retroviruses promote prion-like spreading of proteopathic seeds

**DOI:** 10.1101/2022.05.06.490866

**Authors:** Shu Liu, Stefanie-Elisabeth Heumüller, André Hossinger, Stephan A. Müller, Oleksandra Buravlova, Stefan F. Lichtenthaler, Philip Denner, Ina M. Vorberg

## Abstract

Endogenous retroviruses, remnants of viral germline infections, make up a substantial proportion of the mammalian genome. While usually epigenetically silenced, retroelements can become upregulated in neurodegenerative diseases associated with protein aggregation, such as amyotrophic lateral sclerosis and tauopathies. Here we demonstrate that spontaneous upregulation of endogenous retrovirus gene expression drastically affects the dissemination of protein aggregates between murine cells in culture. Viral glycoprotein Env mediates membrane association between donor and recipient cells and promotes the intercellular transfer of protein aggregates packaged into extracellular vesicles. Proteopathic seed spreading can be inhibited by neutralizing antibodies targeting Env as well as drugs inhibiting viral protein processing. Importantly, we show that also overexpression of a human endogenous retrovirus Env elevates intercellular spreading of pathological Tau. Our data highlight the potential influence of endogenous retroviral proteins on protein misfolding diseases and suggest that antiviral drugs could represent promising candidates for inhibiting protein aggregate spreading.

## Introduction

Neurodegenerative diseases are associated with the aberrant folding of host-encoded proteins into insoluble, highly structured beta sheet-rich protein complexes, termed amyloid. Misfolding of proteins such as the microtubule-binding Tau is associated with highly prevalent Alzheimer’s disease (AD) and other tauopathies. In AD, Tau deposition precedes grey matter atrophy, arguing that aberrant Tau is a major driver of pathogenesis (1). Several different proteins such as TDP-43 or FUS accumulate in the central nervous systems of patients suffering from amyotrophic lateral sclerosis (ALS) or frontotemporal lobar degeneration (FTLD) (2). While mutations in aggregation-prone proteins account for few cases of familial neurodegenerative diseases, the etiologies of spontaneous disease are unknown (3). Protein misfolding occurs through a process of templated conversion, in which small oligomers of misfolded proteins eventually fold into amyloid fibrils capable of templating their aberrant fold onto soluble homotypic proteins. Protein aggregation appears to proceed along neuroanatomical projections, suggesting that intercellular dissemination and propagation of protein misfolding underlies disease progression (4, 5). This process resembles the spreading of prions, infectious protein aggregates composed of PrP that are the causative agents of transmissible spongiform encephalopathies (TSEs) (6). Indeed, a growing number of studies, using cellular models or animals, provide substantial evidence for the spreading of protein aggregates between cells and within tissues (7, 8). Small seeds of aggregated proteins can be either directly released by affected cells or transmitted to bystander cells via direct cell contact (9). Proteopathic seeds capable of inducing protein aggregation in recipient cells can also be packaged into extracellular vesicles (EVs), which are normally secreted by cells for intercellular communication.

We have recently shown that viral glycoproteins such as the vesicular stomatitis virus protein G or SARS Cov-2 spike S expressed by protein aggregate-bearing cells can mediate efficient intercellular contact with bystander cells, resulting in protein aggregate induction in the latter (10). Moreover, viral glycoprotein decoration of EVs from donor cells harboring proteopathic seeds composed of Tau, the yeast Sup35 prion domain NM or prion protein PrP, strongly increased their aggregate inducing capacity in recipient cells. Thus, viral glycoproteins expressed during infection could act as “address codes” that enable delivery, receptor binding, efficient uptake and cytosolic release of proteopathic cargo in recipient cells.

Viral genes are not only encoded by exogenous viruses invading mammalian cells, but are also remnants of mammalian germline infections that happened millions of years ago. Approximately 8-10 % of human and mouse genomes consists of retroviral elements (11). Endogenous retroviruses (ERVs) share a common genome architecture with their exogenous counterparts, in which the coding regions for the capsid proteins (*gag*), reverse transcriptase, integrase and protease (*pol*) and envelope glycoprotein (*env*) are flanked by long terminal repeats. The majority of ERVs are reduced to proviral fragments, and only few are intact or at least contain full open reading frames that are transcribed and/or translated (12). ERVs are subject to tight control by epigenetic modifications that repress transcription (13). Failure to silence ERVs is associated with cancer as well as autoimmune, inflammatory and neurodegenerative disorders (14, 15). Importantly, several human ERV members are upregulated in the brains of tauopathy and ALS patients (16–22). No HERV-derived infectious virions have so far been detected, but HERV-expressing human cell lines can produce viral-like particles (23).

Murine ERV of the Moloney leukemia virus (MLV) clade have integrated into the germline of ancestors millions of years ago. Some inbred mouse lines constitutively generate infectious MLV particles (24). In most inbred mouse lines, however, the propensity of individual MLV loci to produce infectious virions is low. Still, restoration of ERV infectivity by recombination events between distinct loci in immunocompromised mice can result in ERV viremia (25, 26). Here we uncover that de-repression of endogenous MLVs strongly affects spreading of proteopathic seeds between cells. By studying the spreading behavior of cytosolic protein aggregates composed of a yeast prion domain in cell culture, we demonstrate that the reactivation of MLVs strongly increases intercellular aggregate transmission. Reconstitution of HEK donor cells propagating cytosolic NM prions or Tau aggregates with MLV Env was sufficient to promote protein aggregate transfer between cells. Targeting of receptor binding or viral glycoprotein maturation drastically reduced intercellular proteopathic seed spreading. Further, expression of a human ERV glycoprotein also increased Tau aggregate dissemination in cocultures. These findings raise the possibility that de-repression of ERVs accelerates prion-like spreading of protein aggregates and suggests that ERVs represent potential therapeutic targets for disease intervention.

## Results

### Upregulation of endogenous retroviruses correlates with increased aggregate inducing capacity in recipient cells

For this study, we made use of our cell model which is based on the Sup35 prion domain NM. NM is the prion domain of the *Saccharomyces cerevisiae* translation termination factor Sup35 that can form self-templating protein aggregates. The prion domain exhibits compositional similarity to prion-like domains of RNA-binding proteins FUS and TDP-43, known to form protein aggregates in ALS and FTLD (27). Soluble NM expressed in mammalian cells can be induced to aggregate by recombinant NM amyloid fibrils, resulting in cell populations that faithfully replicate NM aggregates over multiple passages (28). Once induced, NM aggregates can also be transmitted to bystander cells by direct cell contact or via EVs, thereby inducing ongoing NM aggregation (29, 30). Using our mouse neuroblastoma N2a Sup35 NM model system, we isolated subclone s2E with HA epitope-tagged Sup35 NM prion aggregates (NM-HA^agg^) (**Suppl. Figure 1a**), which outcompetes other clones in its aggregate inducing capacity in recipient cells (30). For simplicity, we here call this donor clone N2a NM-HA^agg^. Surprisingly, its aggregate-inducing activity in coculture experiments was strongly increased when cells were cultured over prolonged periods of time (**Fig. 1a-c; Suppl. Figure 1b**). The increase in aggregate inducing capacity was reproducible, occurring approximately between 7 to 16 passages post thawing of the cryopreserved cells (approx. 32 to 72 d). Donor cells at high passage number retained their NM aggregate inducing activity even when cryopreserved at passage 21 and subsequently taken into culture (**Suppl. Figure 1c**). For convenience, donor cells cryopreserved at P1 or P21 were subsequently used for experiments for up to 6 passages if not otherwise noted. We refer to cell populations as early and late passage donors (EP and LP, respectively). Increased donor passage number also increased aggregate induction in recipients cultured with conditioned medium from donors. This effect was abolished when conditioned medium was sonicated prior to addition to recipients, suggesting that aggregates might have been contained in EVs that were destroyed by sonication (**Suppl. Figure 1d, e**).

**Figure 1.**
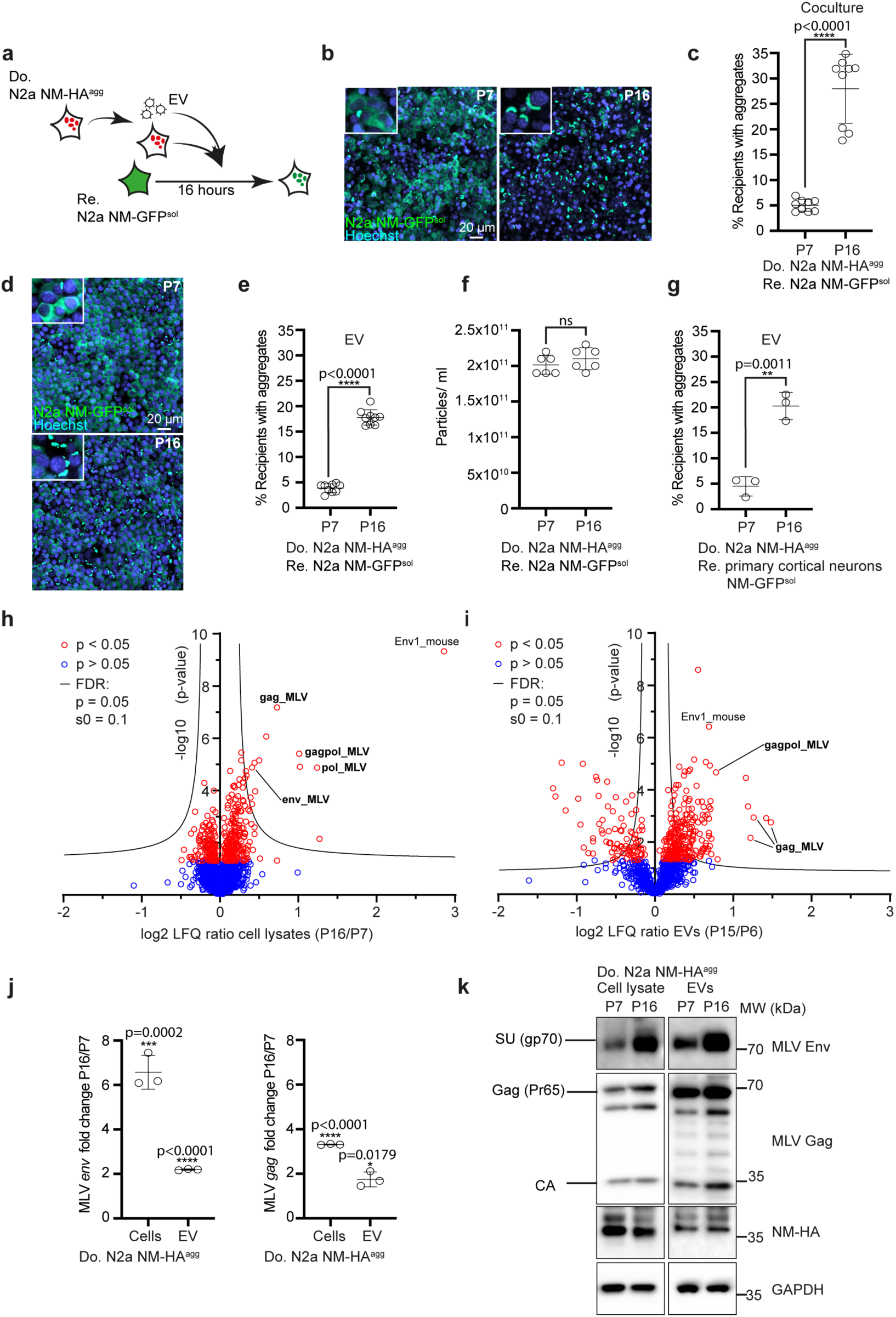
Upregulation of murine endogenous retrovirus in donor cells increases intercellular aggregate induction. **a.** Experimental workflow. Donor N2a NM-HA^agg^ clone passage 6 (P6) and 17 (P17) or EVs derived from the donor clone were added to recipient cells. Automated image acquisition and analysis was performed 16 h post exposure. **b.** Confocal images of cocultures of recipient cells and donor cells P6 or P17. Shown are z stacks. Insets show magnifications of cells. Nuclei were stained with Hoechst. Note that donors have not been stained for NM-HA. **c.** Percentage of recipient cells with induced NM-GFP aggregates upon coculture with donor N2a NM-HA^agg^ cells P6 and P17. **d**. Confocal images of recipient cells cultured in the presence of EV from donors P6 and P17. **e.** Percentage of recipient cells with induced NM-GFP aggregates upon exposure to EVs. **f.** Continuous culture of N2a NM-HA^agg^ cells does not increase EV release. Particles isolated from conditioned medium of cells continuously cultured for 7 (P7) and 16 passages (P16) post thawing were analyzed by ZetaView Nanoparticle Tracking. **g.** Primary neurons expressing soluble NM-GFP were exposed to EVs from donor cells of early or late passage. Quantitative analysis of primary neurons with induced NM-GFP aggregates. **h.** Volcano plot of total cell proteome. Cells lysates of donor N2a NM-HA^agg^ cells at lower and higher passage number were subjected to quantitative mass spectrometry analysis. Proteins were ranked according to their P value and their relative abundance ratio (log2 fold change) in cells of P16 compared to cells of P7. **i.** Volcano plot of proteomes of EVs derived from donor N2a NM-HA^agg^ cells P15 versus P6. Proteins were ranked according to their P value and their relative abundance ratio (log2 fold change) in EVs of P15 compared to EVs of P6. **j.** Continuous culture of N2a NM-HA^agg^ cells increases endogenous *env* and *gag* mRNA as revealed by qRT-PCR. Shown is the fold change in expression in donor cells P16 versus P7. **k.** Increased MLV Env and Gag expression upon continuous cell culture of cells. Shown are Env surface unit SU (gp70) and Gag polyprotein Pr65 and capsid (CA). All data are shown as the means ± SD from three (g, j), six (f) or nine (c, e) replicate cell cultures. Three (c, e-g, j) independent experiments were carried out with similar results. P-values calculated by two-tailed unpaired Student’s t-test. ns: non-significant. Source data are provided as a Source Data file.

We previously demonstrated that NM aggregates are transmitted to bystander cells by EVs. To test if this was also the case at late passage, EVs from donors of different passages were purified by differential centrifugation. Strong aggregate induction was observed with EVs from late passage donors, demonstrating that EVs were involved in cell-free aggregate spreading (**Fig. 1d, e**). Sonication abolished aggregate-inducing activity of EVs (**Suppl. Fig. 1f, g**). Increased aggregate induction was not due to increased EV secretion, as particle numbers did not change significantly over prolonged culture (**Fig. 1f**). EVs isolated from late passage donor cells also increased NM aggregate induction in primary cortical neurons, arguing that the effect was independent of recipient cells (**Fig. 1g**).

To identify changes in the proteome of donor cells that might contribute to protein aggregate spreading, we performed mass spectrometry analyses of total cell lysate (**Fig. 1h**) and donor EV fractions (**Fig. 1i**) at early and late passages. Among the proteins increased in donor cells and EVs upon prolonged culture, we identified mouse endogenous γ-retroviruses MLV proteins to be highly and significantly increased (**Source data)**. Endogenous MLV are remnants of ancient germline infections that constitute approx. 10 % of the mouse genome. Partially overlapping MLV open reading frames code for polyproteins Gag (comprising matrix, p12, capsid and nucelocapsid), Pol (reverse transcriptase, integrase and protease) and for the envelope glycoprotein Env, giving rise to full genome RNAs as well as mRNAs coding for *gag/pol* and *env*, respectively. Prolonged cell culture increased mRNA levels coding for *env* and *gag* in both cell lysates and EVs (**Fig. 1j**). Western blot analyses confirmed increased expression of Env and Gag in cell lysates and EV fractions from donors upon prolonged culture (**Fig. 1k**). Gag was processed into capsid (CA, p30), nucleocapsid core (MA) and nucleocapsid (NC) by the viral protease. Env was cleaved into a surface subunit (SU, gp70) and a transmembrane domain (TM, Pr15/p15E) (31).

### Donor cells produce both active viral particles and EVs

The finding that MLV mRNA and proteins increased in donor cells upon prolonged culture suggested that donor cells secrete active retrovirus, as has been observed for few cell lines before (32–35). In line with this, increased reverse transcriptase activity was observed upon prolonged culture of donors (**Fig. 2a**). MLVs are classified by their respective Env proteins as ecotropic, polytropic and xenotropic, depending on their receptor preferences (36, 37). To test if expression of ERVs had been triggered by NM aggregation, we preformed quantitative real-time PCR on mRNA extracted from different N2a NM populations before and after exposure to recombinant NM fibrils. Aggregate induction had no influence on induction of endogenous MLV subgroups, arguing that upregulation of MLV is likely influenced by other means (**Suppl. figure 2a-f**).

**Figure 2.**
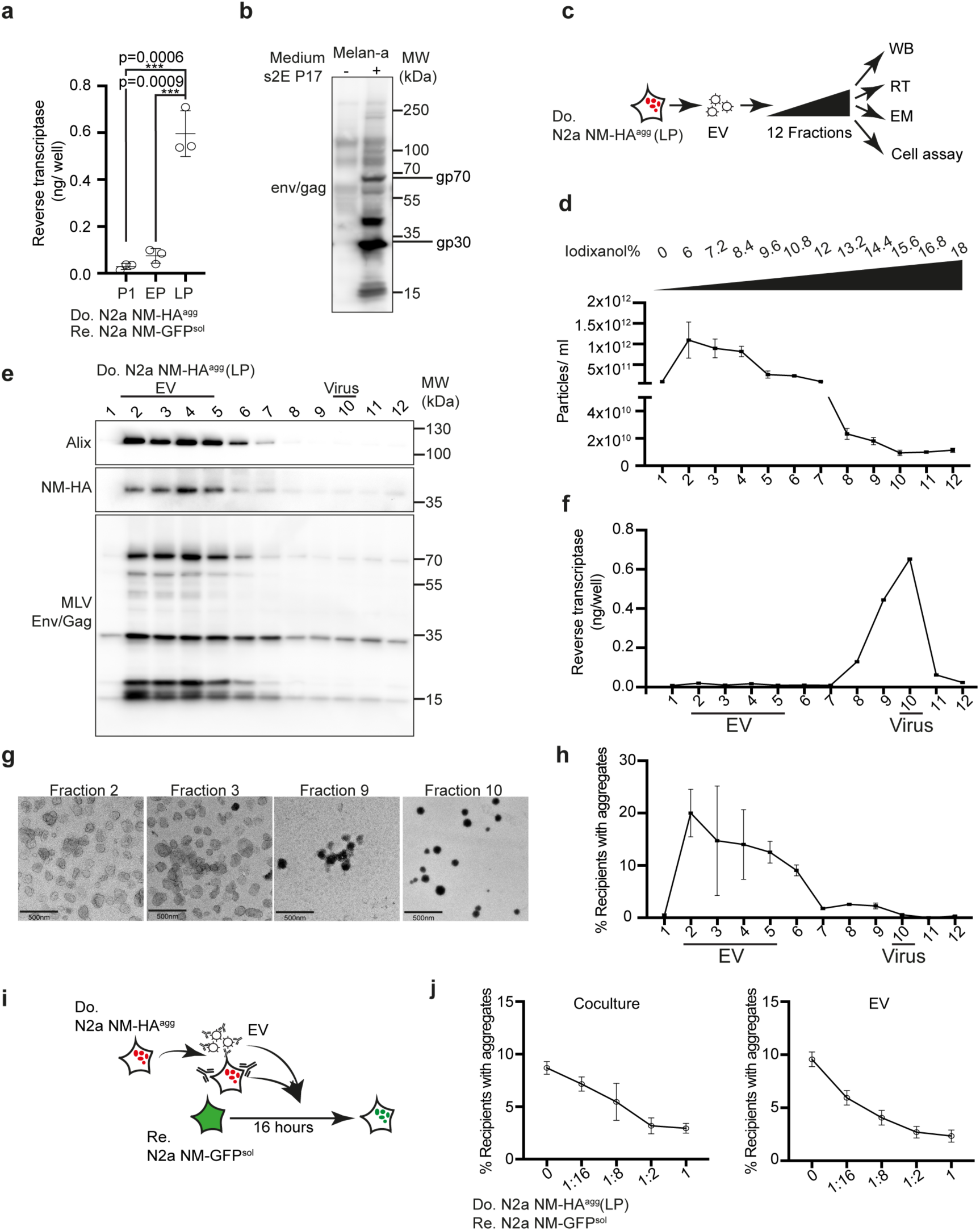
Donor cells of late passage produce both EVs and active retroviral particles. **a.** Viral particles and EVs were precipitated from conditioned medium of N2a NM-HA^agg^ cells of different passage numbers using polyethylene glycol. Reverse transcriptase (RT) activity was determined using a colorimetric RT assay (Roche). **b.** Viral particles released from donor N2a NM-HA^agg^ cells at late passage are infectious to a murine melanocytic cell line susceptible to MLV infection (57). Melan-a cells were exposed to conditioned medium of the donor clone. Western blot analysis was performed 6 d later using antibody ABIN457298 against xenotropic MLV viruses detects expression of Env and Gag only in infected melan-a cells. Viral proteins are indicated. **c.** Experimental workflow. The 100,000 x g pellet from conditioned medium of donor clone N2a NM-HA^agg^ (late passage) was subjected to a shallow OptiPrep density gradient. Individual fractions were analyzed for particle numbers, reverse transcriptase (RT) activity, particle morphology by transmission electron microscopy (TEM), protein content by Western blot (WB) and aggregate inducing activity. **d.** Particle numbers of gradient fractions determined using ZetaView. **e.** Density gradient fractions were analyzed for Alix, endogenous Env/Gag (ABIN457298) and NM-HA by Western blot. **f.** Reverse transcriptase (RT) activity in individual fractions is used to identify viral particles (fractions 8-11). **g.** Transmission electron microscopy reveals the typical cup-shaped EVs in fractions 2 and 3 and virus particles with an electron dense core in fractions 9 and 10. Scale bar: 500 nm. **h.** NM-GFP aggregate inducing activity of gradient fractions in recipient cells. N2a NM-GFP^sol^ cells exposed to different fractions were analyzed for induced NM-GFP aggregates 16 h post exposure. **i.** Experimental workflow. Recipient cells (LP) were cocultured with donor cells in the presence of different dilutions of anti-MLV env antibody mAb83A25 for 1 h. Alternatively, recipients were cultured with donor EVs that had been pre-incubated with anti-env antibodies for 1 h. Anti-env antibodies were present throughout the experiment. The percentage of recipient cells with induced NM-GFP aggregates was determined 16 h post coculture or EVs addition. All data are shown as the means ± SD from three (a, d, i) or twelve (j) replicate cell cultures. Three (a, d, i, j) independent experiments were carried out with similar results. P-values calculated by two-tailed unpaired Student’s t-test (a). Source data are provided as a Source Data file.

To demonstrate that donor cells produced infectious virus, vesicle fractions derived from donors were added to murine melan-a cells, a cell line permissive for MLV. Detection of Gag and Env by Western blot demonstrated that cells produced infectious virus (**Fig. 2b**) (38). We further tested if Vectofusin-1, a compound that increases viral interaction with cellular membranes, could enhance aggregate induction. Aggregate induction was also increased when conditioned medium was added to recipients in the presence of Vectofusin-1 (**Suppl figure 2g, h**) (39). To test which vesicle fractions released by the donor cells were NM-seeding competent, we separated EVs from viral particles by an OptiPrep velocity gradient previously used to separate HIV-1 virions from microvesicles (**Fig. 2c**) (40). Fractions with highest particle concentrations (fractions 2-6) harbored Alix and NM-HA, arguing that they contained EVs (**Fig. 2d, e**). Gag and Env were distributed throughout the gradient, with highest levels present in Alix-positive fractions (**Fig. 2e**). Reverse transcriptase (RT) activity was almost exclusively present in fractions 8-11, arguing that these fractions contained active virus (**Fig. 2f**). This was confirmed by electron microscopy, which revealed membranous 80-100 nm spherical particles with an electron-dense core, characteristic of γ-retroviral particles in fractions 9 and 10 (**Fig. 2g**) (41). By contrast, vesicles in fractions 2 and 3 exhibited a cup-shaped morphology, characteristic of EVs in TEM (42). Next, we tested the NM aggregate inducing activity of fractions by adding them to recipient N2a NM-GFP^sol^ cells (**Fig. 2h**). Highest aggregate induction was associated with RT-negative EV fractions 2-6. Interestingly, aggregate induction could also be inhibited by antibodies against MLV Env in both coculture and by EVs (**Fig. 2i**). We conclude that NM aggregate seeding activity is mainly associated with the RT-negative EV fraction. Still, aggregate induction can be inhibited by antibodies directed against MLV Env.

### Modulation of endogenous retrovirus expression affects intercellular NM aggregate induction

The foregoing experiments suggested that Env plays a prominent role in intercellular aggregate transmission and induction. We tested if silencing of MLV in donors affects aggregate induction in recipient cells. Transfection of donor cells with three individual siRNAs targeting MLV *env* or *gag/pol* only slightly decreased MLV gene products, likely due to multicopy ERVs (24, 43) (**Fig. 3a-c**). Still, knock-down of either *gag/pol* or *env* significantly decreased NM aggregate induction in cocultured N2a NM-GFP cells (**Fig. 3d**). Since Gag/Pol and Env are encoded by different mRNAs, these results argue that Env as well as Gag/Pol contribute to enhanced proteopathic seed spreading.

**Figure 3.**
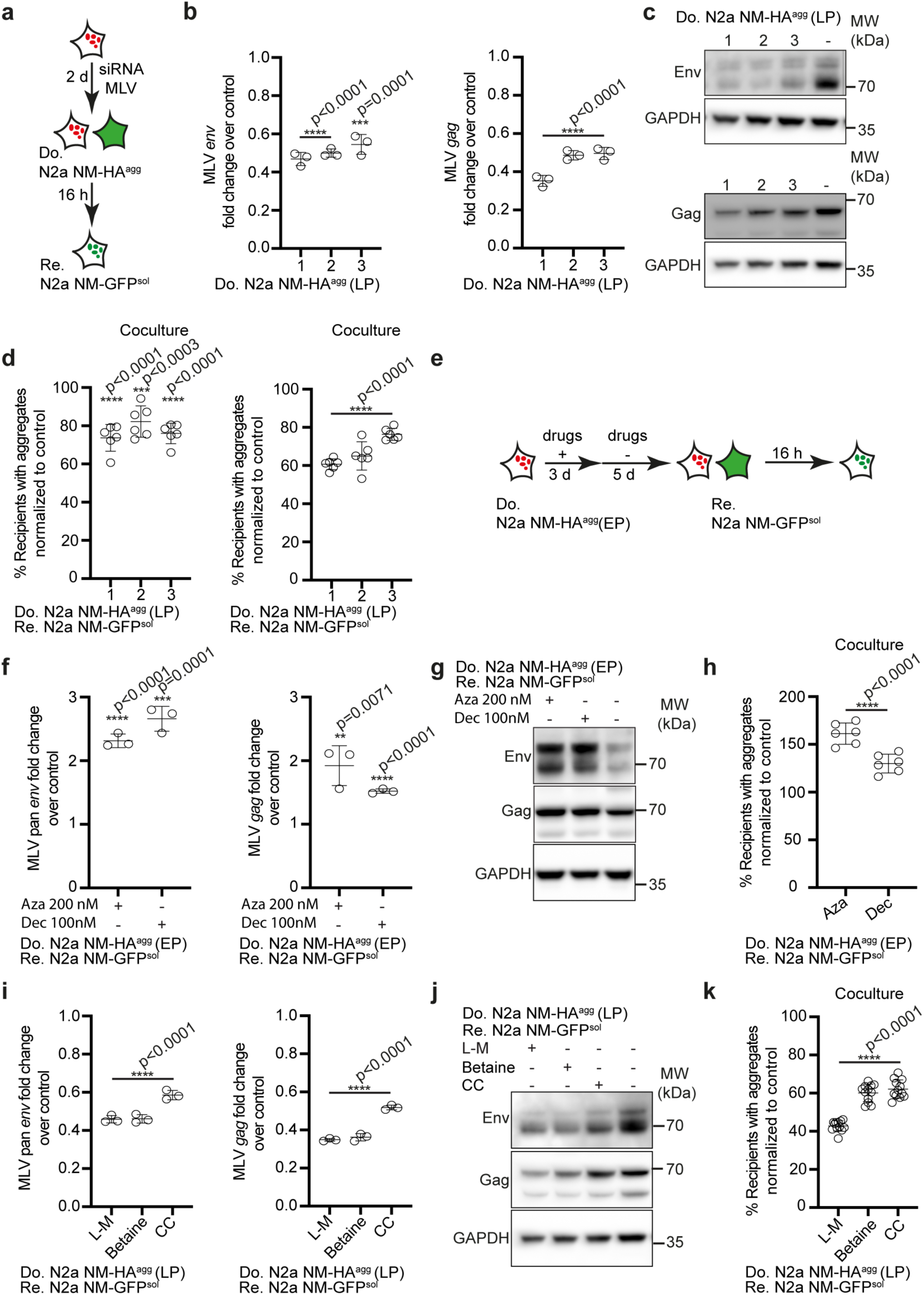
Epigenetic modulation of MLV expression affects aggregate spreading. **a.** Experimental scheme. Donor N2a NM-HA^agg^ cells of late passage was transfected with three individual siRNAs against endogenous *env* or *gag* and non-silencing siRNA as control. Donors were subsequently cocultured with recipients. **b.** Reduction of donor transcripts following siRNA treatment assessed by qRT-PCR. Shown are fold changes relative to mock control. **c.** Western blots of Env and Gag expression in N2a NM-HA^agg^ cells (LP) transfected with three anti-*env* or anti-*gag* siRNAs or control siRNA. GAPDH serves as a loading control. **d.** Donors cocultured with recipient N2a NM-GFP^sol^ cells 72 h post transfection. Shown is the percentage of recipient cells with NM-GFP^agg^ normalized to control**. e.** Coculture experiments with methyltransferase inhibitors. The donor clone (EP) was treated with DNA methyltransferase inhibitors 5-azacytidine (Aza), 5-aza-2’- deoxycytidine (Dec) or DMSO for 3 d and then cultivated in normal medium for 5 d. Cells were subsequently cocultured with recipients for 16 h. **f.** Expression of MLV *env* (left panel) or *gag* (right panel) transcripts in donor cells 5 d post treatment was assessed by qRT-PCR. **g.** Western blot of Env and Gag upon treatment with methyltransferase inhibitors. GAPDH served as a loading control. **h.** Percentage of NM-GFP^agg^ positive recipient cells cocultured with methyltransferase inhibitor-treated donors normalized to control coculture with solvent-treated donors. **i.** Donor cells of high passage number (LP) were treated with methyl group donors L-methionine (L-M), betaine, choline chloride (CC) or medium control for 6 d. MLV *env* or *gag* transcripts were analyzed by qRT-PCR. **j.** Expression of Env and Gag upon treatment with methyl group donors for 6 d. GAPDH served as a loading control. **k.** Percentage of NM-GFP^agg^ positive recipient cells cocultured with drug-treated donors normalized to solvent-treated donors. All data are shown as the means ± SD from three (b, f, i), six (d, h) or twelve (k) replicate cell cultures. Three (b, d, f, h-i, k) independent experiments were carried out with similar results. P-values calculated by two-tailed unpaired Student’s t-test. Source data are provided as a Source Data file.

Expression of MLV underlies epigenetic control by silencing of proviral promoter regions through DNA methylation early during development (44). As DNA methyltransferase inhibitors can induce expression of ERVs (45), we tested if erasing epigenetic marks affects aggregate inducing activity of donor cells (**Fig. 3e**). Treatment of donor cells at early passage with DNA methyltransferase inhibitors 5-Azacytidine (Aza) or Decitabine (Dec) (46, 47) resulted in increased expression of total MLV *env* and *gag* mRNA (**Fig. 3f**) and increased Env and Gag protein (**Fig. 3g**). Both drugs also significantly increased NM aggregate induction in recipient cells when these were cocultured with pretreated donors (**Fig. 3h**). In a reverse experiment, increased methylation with treatment of late passage donors with l-methionine, betaine or choline chloride resulted in decreased *env* and *gag/pol* mRNA (**Fig. 3i**), protein levels (**Fig. 3j**) and reduced aggregate induction in recipients (**Fig. 3k**). The findings that siRNA against MLV mRNAs and epigenetic drugs that modulate MLV expression affect intercellular aggregate spreading suggest that MLV expression in involved intercellular aggregate spreading in our N2a cell culture system.

### Inhibition of MLV protein maturation impairs intercellular NM aggregate induction

MLV proteins Env and Gag require the MLV viral protease for processing into mature proteins. (48–50). To investigate if EV-mediated NM aggregate induction depends on proper maturation of viral proteins, we tested anti-HIV-1 drugs for their effects on NM aggregate induction in coculture. Among the four tested HIV protease inhibitors, Amprenavir and Atazanavir have previously been shown to inhibit MLV protease (51). Treatment of cocultures with these compounds (**Fig. 4a**) had no effect on the percentage of donor cells with pre-existing NM-HA aggregates (**Suppl. figure 3a**). However, the two MLV protease inhibitors Atazanavir and Amprenavir reduced the percentage of recipient cells with induced NM-GFP aggregates (**Fig. 4b**). Other HIV protease inhibitors, reverse transcriptase inhibitors or Hepatitis C virus (HCV) protease inhibitors had no effect on donor aggregates or aggregate induction in recipients in coculture (**Suppl. figure 3b**).

**Figure 4.**
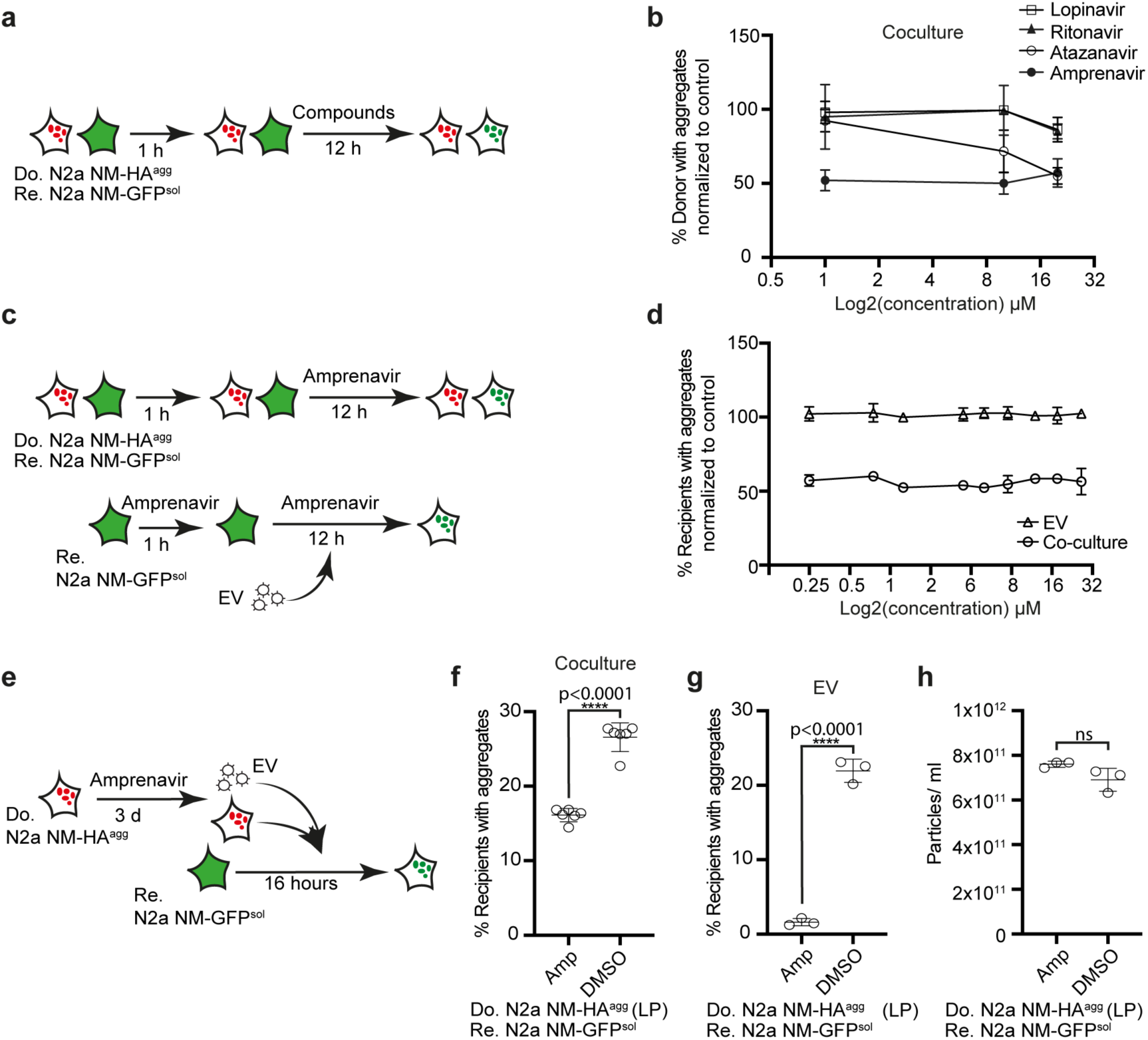
Protease inhibitors blocking MLV maturation impair intercellular protein aggregate spreading. **a.** Workflow of compound test in cocultures. Donor clone N2a NM- HA^agg^ (LP) and recipient N2a NM-GFP^sol^ cells were co-seeded and viral protease inhibitors were added at three concentrations 1 h later. 12 h post drug treatment, donor and recipient cells were analyzed for the percentage of donor cells with NM-HA^agg^ or recipient cells with induced NM-GFP^agg^. **b.** Quantitative analysis of the effect of protease inhibitors on cells with NM-GFP aggregates. Recipients with NM-GFP^agg^ that were solvent-treated (DMSO) were set to 100 %. **c.** Workflow of compound test in cocultures. Donor N2a NM-HA^agg^ (LP) and recipient N2a NM-GFP^sol^ were co-seeded and exposed to different concentrations of Amprenavir 1 h later. Alternatively, recipient N2a NM-GFP^sol^ cells were pretreated with different concentrations of Amprenavir for 1 h, and cells were subsequently exposed to donor-derived EVs for 12 h. Note that donor cells (LP) from which EVs were isolated remained untreated. **d.** Amprenavir effects on percentage of recipient cells with induced NM-GFP aggregates in the two assays. Results were normalized to solvent control. **e.** Experimental procedure. Donor N2a NM-HA^agg^ cells (LP) were treated with Amprenavir for 3 d. Cells were subsequently cocultured with recipient N2a NM-GFP^sol^. Alternatively, EVs from treated donors (LP) were added to recipient cells. Aggregate formation in recipients was assessed 16 h later. **f.** Amprenavir inhibits NM-GFP aggregation in cocultured recipient cells. **g.** Effect of Amprenavir-treatment of donor cells (LP) on the induction of NM-GFP aggregates in recipient cells upon exposure to donor-derived NM- HA^agg^ bearing EVs. **h.** Effect of Amprenavir on particle release by donor cells (LP). All data are shown as the means ± SD from three (g-h) or six (f) replicate cell cultures. Three (f-h) independent experiments were carried out with similar results. P-values calculated by two-tailed unpaired Student’s t-test. ns: non-significant. Source data are provided as a Source Data file.

We hypothesized that Amprenavir inhibited MLV protease-driven viral protein maturation in the donor cells or during EV formation and thus had no effect on recipients. Thus, we tested the effect of the HIV protease inhibitor Amprenavir on NM-GFP aggregate induction by treating recipient cells 1 h prior to addition of EVs isolated from donor cells (LP) and incubating cells in the presence of drug for further 12 h. As a control, donor cells (LP) were also cocultured with recipients in the presence of the drug (**Fig. 4c**). As expected, Amprenavir only inhibited aggregate induction in cocultures, but not in drug-treated recipients exposed to EVs (**Fig. 4d**). In a reverse experiment, Amprenavir pretreated donor cells (LP) were cocultured with recipient cells in the absence of Amprenavir. Additionally, EVs isolated from Amprenavir-treated donors (LP) were tested for their aggregate inducing activity in recipient cells (**Fig. 4e**). Amprenavir pre-treatment of donor cells significantly inhibited intercellular aggregate induction during coculture (**Fig. 4f**) and also basically abolished EV-mediated aggregate induction in recipient cells (**Fig. 4g**), without affecting secreted particle numbers (**Fig. 4h**). These experiments suggest that maturation of endogenous MLV encoded gene products in donor cells or donor-derived EVs is required for efficient aggregate induction in recipient cells.

### Viral Envelope binding to its receptor XPR1 is required for intercellular aggregate induction

MLV envelope proteins mediate specific contact of virions with their receptors on target cells and induce cargo release into the cytosol by enforcing fusion of lipid bilayers. According to our proteomic analysis, polytropic *env* was upregulated in cells of late passage, suggesting that contact between polytropic Env and its receptor XPR1 was involved in NM aggregate spreading. Silencing of recipient MLV receptor XPR1 but not mCat-1, the receptor for ecotropic MLV (**Fig. 5a, b**), strongly reduced NM aggregate induction in cocultures (**Fig. 5c**) and by EV exposure (**Fig. 5d**), confirming that upregulated MLVs belong to the group X/P-MLVs using this receptor. Silencing of both receptors in recipient cells had no effect on NM aggregate induction by recombinant NM fibrils, arguing that NM aggregate uptake was mediated by EV-receptor contact and not the direct contact of an NM seed with the receptor (**Suppl figure 4a**).

**Figure 5.**
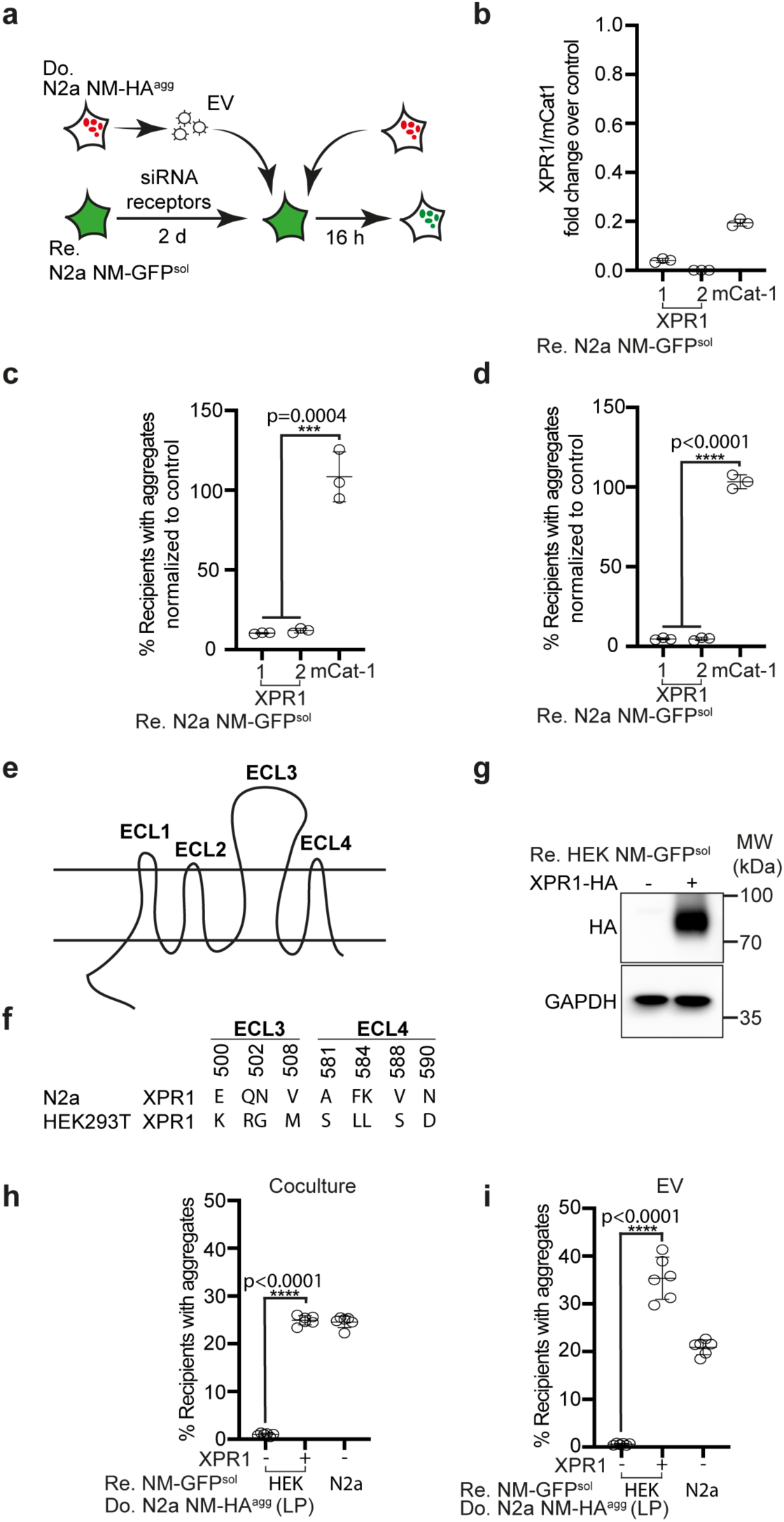
Receptor polymorphisms modulate intercellular proteopathic seed spreading. **a.** Experimental workflow. Recipient N2a NM-GFP^sol^ cells were transfected with two siRNAs against XPR1, one mCat-1 siRNA or a non-silencing siRNA control (mock). 48 h later, recipient cells were cocultured with donor cells (LP). Alternatively, recipients were exposed to EVs isolated from conditioned medium of N2a NM-HA^agg^ cells (LP). NM-GFP aggregate induction was determined 16 h post exposure or coculture. **b.** Knock-down of receptor mRNA in N2a NM-GFP^sol^ cells was assessed 48 h post siRNA transfection by qRT-PCR. Shown is the fold change of mRNA expression normalized to mock control. **c., d.** Recipients with NM-GFP aggregates following receptor knock-down. Shown are results of coculture (**c**) recipients exposed to donor derived EVs (LP) (**d**). NM-GFP aggregate induction was measured 16 h post fibril/EV addition or coculture. **e.** Transmembrane structure of XPR1. The receptor contains 4 extracellular loops (ECL1-4) (37). **f.** Polymorphic variants of xenotropic and polytropic (X/P-) MLV receptor XPR1 in mouse N2a and human HEK cells. Shown are mismatches in the surface-exposed loops ECL 3 and 4. ECL3 and 4 are required for binding of X/P-MLV (37). **g.** Ectopic expression of the N2a XPR1 receptor variant in poorly permissive HEK NM-GFP^sol^ cells. Ectopically expressed XPR1-HA was detected using anti-HA antibodies. GAPDH served as a loading control. **h.** HEK NM-GFP^sol^ cells were transfected with mouse XPR1-HA or mock transfected and subsequently cocultured with donor N2a NM-HA^agg^ cells (LP). **i.** Alternatively, transfected HEK NM-GFP^sol^ cells were exposed to EVs from N2a NM- HA^agg^ cells (LP). All data are shown as the means ± SD from three (b-d) or six (h-i) replicate cell cultures. Three (b-d, h-i) independent experiments were carried out with similar results. P-values calculated by two-tailed unpaired Student’s t-test. Source data are provided as a Source Data file.

XPR1 is a receptor with eight putative transmembrane domains and four extracellular loops (ECL) (**Fig. 5e**) (52). At least six polymorphic variants of XPR1 restrict infection by specific X/P-MLVs, with polymorphisms in ECL 3 and 4 affecting entry of certain X/P-MLV subtypes (53). Analysis of XPR1 of susceptible N2a NM-GFP^sol^ cells demonstrated that its Env recognition domain differed at 9 residues within ECL 3 and 4 from XPR1 expressed by HEK NM-GFP^sol^ cells, a cell line refractory to NM aggregate induction by N2a NM-HA^agg^-derived EVs (**Fig. 5f**). Expression of the N2a XPR1 variant (**Fig. 5g**) conferred susceptibility to HEK cells, both in coculture or upon exposure to donor-derived EVs (**Fig. 5h, i**). By contrast, expression of the N2a polymorphic XPR1 variant had no effect on aggregate induction by recombinant NM fibrils (**Suppl fig. 4b**). We conclude that efficient NM aggregate induction via coculture or EVs depends on specific interaction of Env with its receptor.

### Retroviral protein expression and viral particle formation promote intercellular spreading of NM and Tau aggregates

The foregoing experiments demonstrated that epigenetic upregulation of MLV strongly affected intercellular aggregate spreading. So far it was unclear if generation of active viral particles was required for this process. To test this, HEK NM-GFP^agg^ cells not coding for MLV were transfected with combinations of plasmids coding for MLV *gag/pol*, amphotropic MLV *env* 10A1 and MLV transfer vector (coding for mCherry) for virus production (**Fig. 6a**). Vero cells stably expressing NM-GFP^sol^ (10) were chosen as recipients due to their high expression level of amphotropic Env receptor Pit-2 (**Fig. 6b**). Ectopic expression of viral proteins Env and Gag/Pol resulted in significantly increased aggregate induction rates in cocultured recipients, with higher induction rates when *env*, and *gag/pol* were expressed simultaneously (**Fig. 6c, d**). Interestingly, highest induction rates were observed when donor cells also produced virus (**Fig. 6e, f**).

**Figure 6.**
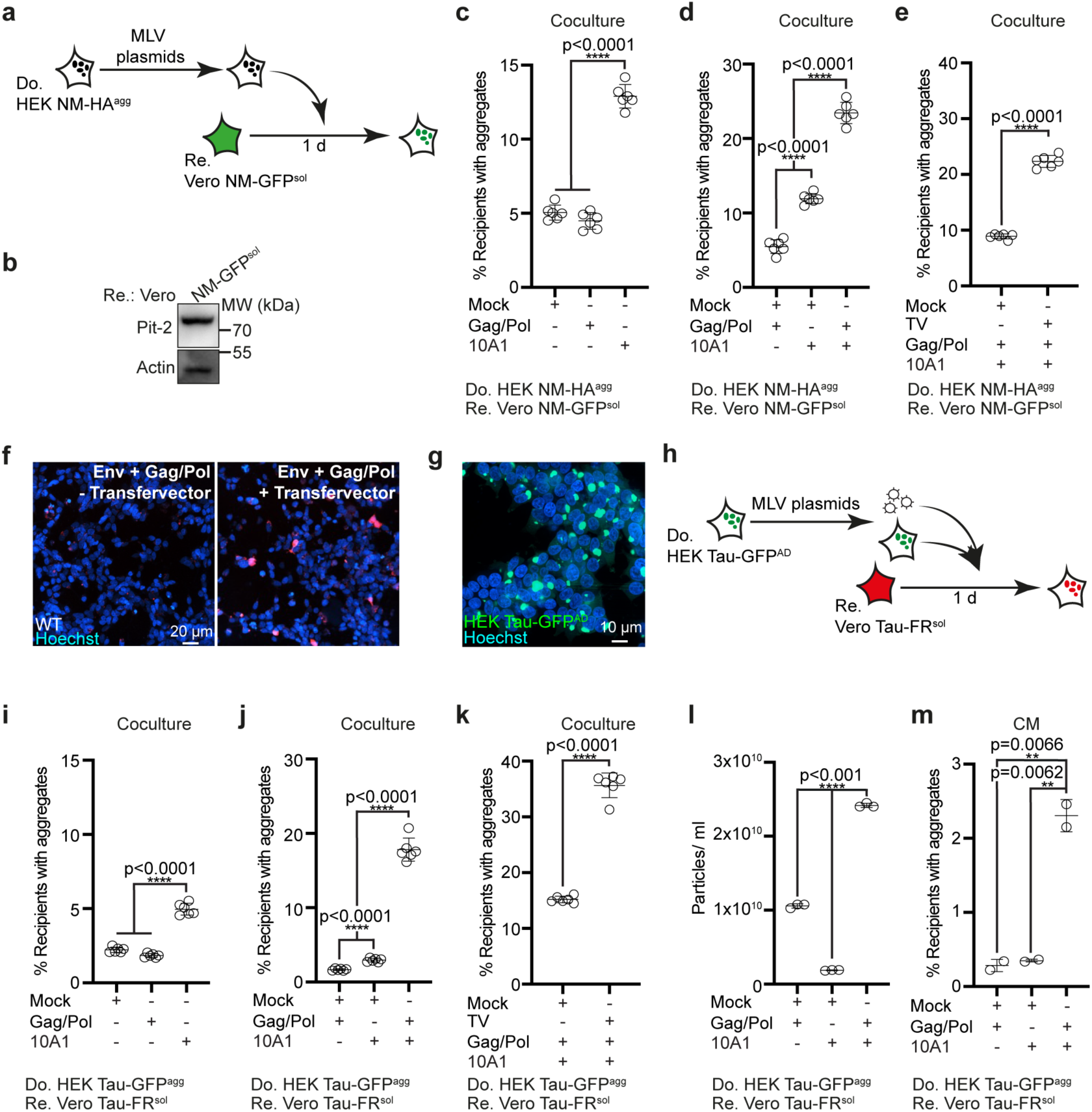
Impact of MLV reconstitution on intercellular NM and Tau aggregate induction. **a.** Schematic overview. HEK NM-HA^agg^ donor cells were transfected with combinations of for amphotropic MLV *env* 10A1 vector, *gag/pol* plasmid, retroviral transfer vector and/or empty vector. Donors were subsequently cocultured with recipient Vero NM-GFP^sol^ cells. **b.** Western blot analysis of cell lysates from recipient cells expressing Pit-2. **c.** Percentage of recipient Vero cells with induced NM-GFP^agg^ cocultured with donors transfected with single plasmids. **d.** Percentage of recipient cells with induced NM-GFP^agg^ cocultured with donors transfected with combinations of plasmids coding for *env*, *gag/pol* or non-viral empty plasmid. **e.** Percentage of recipient cells with induced NM- GFP^agg^ cocultured with donors that were additionally transfected with/without transfer vector (TV) for virus production. **f.** Donor cells transfected with/ without mCherry-coding retroviral transfer vector and plasmids coding for *gag/pol* and *env* produce virus that is infectious to wildtype cells. **g.** Representative image of HEK Tau-GFP^AD^ donor cell population. **h.** HEK Tau-GFP^AD^ donor cells were transfected with combinations of plasmids coding for amphotropic MLV *env* 10A1, *gag/pol*, retroviral transfer vector and/or non-viral empty vector. Donors were subsequently cocultured with recipient Vero Tau- FR^sol^ cells. Alternatively, donor EVs were added to recipient cells. **i.** Confocal image of HEK Tau-GFP^AD^ donor cells. **i.** Percentage of recipient cells with induced Tau-FR^agg^ upon coculture with donors transfected with individual plasmids. **j.** Percentage of recipient cells with induced Tau-FR^agg^ upon coculture with donors transfected with plasmid combinations. **k.** Percentage of recipient cells with induced Tau-FR^agg^ upon coculture with donors transfected with plasmid combinations and transfer vector. **l.** Transfection of plasmids coding for *gag/pol* and *env* simultaneously increases particle secretion. **m.** Percentage of recipient cells with Tau-FR^agg^ exposed to conditioned medium of donors adjusted for comparable particle numbers. All data are shown as the means ± SD from two (m), three (l) or six (c-e, i-k) replicate cell cultures. Three (c-e, i-m) independent experiments were carried out with similar results. P-values calculated by two-tailed unpaired Student’s t-test (e, k-m) or one-way ANOVA. (c, d, i, j, l). Source data are provided as a Source Data file.

Next, we tested if retroviral proteins also promoted intercellular transmission of protein aggregates associated with neurodegenerative diseases. To this end, we made use of our recently developed cell culture model propagating aggregates composed of the repeat domain of a mutant human Tau protein (**Fig. 6g**) (54). HEK cells stably expressing a soluble GFP-tagged repeat domain variant of human Tau (hereafter termed Tau-GFP^sol^) stably produce and maintain Tau-GFP aggregates upon exposure to Alzheimer’s disease brain homogenate (Tau-GFP^AD^) (10, 54). Upon donor transfection with retroviral plasmids (**Fig. 6h**), induction rates in cocultured Vero cells expressing the same Tau variant fused to FusionRed (Tau-FR^sol^) (10) were significantly increased (**Fig. i-k**). Again, highest induction rates were observed when donors were transfected with plasmids coding for *gag/pol, env* and the transfer vector (**Fig. 6k**). Transfection of combinations of *gag/pol* and *env* vectors also strongly increased vesicle release (**Fig. 6l**). When conditioned medium was adjusted for comparable particle numbers, highest induction rates were observed when donors were transfected with both *gag/pol* and *env* constructs (**Fig. 6m**). We conclude that expression of retroviral proteins Env, Gag and Pol is sufficient to promote intercellular proteopathic seed spreading, but that proteopathic seed spreading is most efficient when donor cells produce active viral particles.

### Human endogenous retrovirus Env can increase proteopathic seed spreading

In contrast to murine endogenous retroviruses of the MLV clade, HERV have so far not been shown to produce infectious virions in vivo. However, under certain circumstances, HERVs become reactivated, resulting in transcript and even protein expression. We thus tested if Env proteins encoded by HERVs could affect spreading of Tau misfolding. To this end, donor HEK Tau-GFP^AD^ cells were transfected with a plasmid coding for the HERV-W Env Syncytin-1 or empty vector (**Fig. 7a, b**). Coculture experiments with both HEK and Vero cells expressing Tau-FR^sol^ revealed that Syncytin-1 resulted in a significant increase in recipient cells with Tau-FR aggregates (**Fig. 7c-f**). We conclude that gene products of both murine and human ERVs can facilitate intercellular protein aggregate spreading.

**Figure 7.**
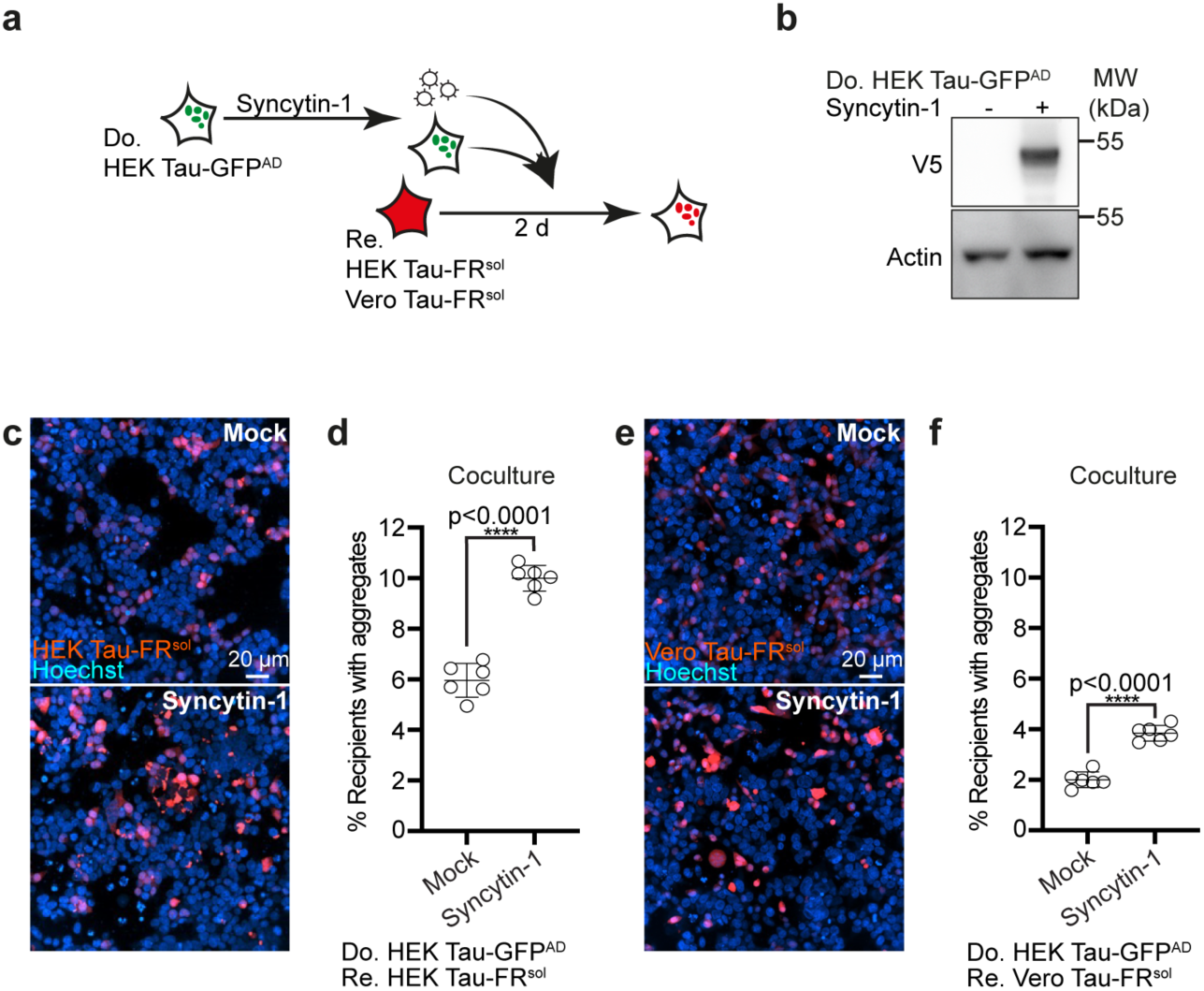
HERV-W Env interactions with its receptors increase Tau aggregate induction. **a.** Experimental workflow. Donor HEK cells stably propagating aggregated Tau-GFP^AD^ were transfected with plasmid coding for V5 epitope-tagged Syncytin-1 (Syn- V5) and were subsequently cocultured with recipient HEK or Vero cells expressing Tau- FR^sol^. **b.** Western blot analysis of donor clone transfected with plasmids coding for Syn- V5. **c.** Coculture of donor and HEK recipient cells. Note that we have not stained the donors in this experiment. **d.** Quantitative analysis of the percentage of recipient cells with induced aggregates upon coculture. **e.** Coculture of donor and Vero recipient cells. **f.** Quantitative analysis of the percentage of recipient cells with induced aggregates. All data are shown as the means ± SD from 6 (d, f) replicate cell cultures. Three (d, f) independent experiments were carried out with similar results. P-values calculated by two-tailed unpaired Student’s t-test. Source data are provided as a Source Data file.

## Discussion

Accumulating evidence argues that ERVs, resulting from retroviral germline invasions throughout evolution, are upregulated in NDs. While some HERV gene products can be directly neurotoxic, such as HERV-W Env (55) and HERV-K Env (56), also inflammatory responses due to viral transcripts have been implicated in ND development (19). Our data suggest an additional mechanism of how ERV proteins could contribute to neurodegeneration, namely by accelerating intercellular dissemination of protein particles. Activation of polytropic endogenous MLV proviruses resulted in the production of infectious virions as well as the secretion of protein aggregate-loaded EVs decorated with viral Env. As a result, MLV upregulation drastically increased the intercellular transmission of proteopathic seeds by EVs to bystander cells or to cells in direct contact and induced protein aggregation in the latter.

Which cellular mechanism accounts for the consistent epigenetic switch upon continuous cell culture remains to be explored. The effect of epigenetic drugs on MLV expression argues that demethylation of MLV promoter regions at least partially explains this phenomenon. The activation of ERV was independent of transgene expression or the induction of protein aggregates. Activation and generation of ERV-derived retroviral particles has been reported for several cell lines in culture, including N2a cells (32-35, 57, 58). Interestingly, receptor usage of viral particles produced in our N2a cell model clearly differed from the ecotropic MLV identified by others, arguing that within a given cell population, different MLVs can become de-repressed (33, 57).

Our detailed analysis reveals that the effect of reactivated endogenous MLV on protein aggregate spreading can be attributed to the expression of Env glycoprotein and retroviral Gag/Pol polyproteins. Upon increased expression, Env on the cell surface or on EVs mediates the binding to specific receptors on the cell surface of recipient cells. Cleavage of the MLV Env R-peptide by MLV protease then mediates fusion of cell membranes or EVs with the recipient cell or its endo-lysosomal membranes (48), resulting in the release of proteopathic seeds into the cytosol of the recipient cell and subsequent aggregate induction. Surprisingly, reconstitution experiments in HEK cells demonstrate that MLV Env alone is sufficient to increase intercellular aggregate spreading, in analogy with our findings that vesicular stomatitis virus G and SARS-CoV2 spike S glycoproteins can elevate intercellular aggregate induction (10). The additional positive effect of co-transfecting a plasmid coding for Gag and Pol polyproteins on aggregate induction is likely due to more efficient R peptide cleavage as well as concentration of Env and Gag polyprotein within rafts (48, 59). However, highest induction rates were achieved when all plasmids for MLV virus production were transfected into donors. Thus, cells harboring protein aggregates and also actively producing infectious MLV virus most efficiently induce proteopathic seeds in bystanders. The reason for this is unclear, but might be related to the fact that MLV RNA increases the efficiency of proper Gag-Gag interactions, which in turn also affect proper Env positioning within rafts (60).

Most experiments in this study were performed using a model protein for cytosolic protein aggregates, which is based on the prion domain of the *S. cerevisiae* translation termination factor Sup35. The prion domain of Sup35 shares striking compositional similarity with so-called prion-like domains of a growing number of proteins associated with ALS and FTLD, such as FUS and TDP-43 (27). In analogy to our study, experiments with transgenic mice have recently demonstrated that misfolded aggregated TDP-43 is secreted within EVs, suggesting that protein aggregates with similar domains are secreted by the same mechanism (61). Further, EVs isolated from plasma of ALS patients are enriched for TDP-43 and FUS (62). Thus, it is feasible to assume that other protein aggregates with prion-like domains might also be affected by ERV de-repression. This hypothesis is supported by our findings that retroviral gene expression in donor cells harboring Tau aggregates also increased aggregate dissemination, demonstrating that this effect was independent of the type of cytosolic protein aggregate. Our results are consistent with a previous study, showing that simultaneous infection of cells with scrapie agent and friend retrovirus strongly enhanced intercellular spreading of pathologic prion protein and scrapie infectivity (63). Viral Env and Gag association with prion-containing EV fractions has previously been observed for a prion-infected, endogenous ecotropic MLV producing N2a subpopulation. However, the role of Env expression in intercellular prion spreading has not been studied (33). Interestingly, independent in vivo co-infections with MLV and scrapie showed no effect on scrapie incubation times, potentially because target cells for exogenous MLV and scrapie differ (64, 65). Unfortunately, the effect of de-repressed MLVs on spreading of proteopathic seeds cannot easily be tested in ND mouse models, as endogenous MLV proviruses in common mouse lines such as C57BL/6 used in ND research are transcribed at low or undetectable levels (43, 66). Endogenous polytropic MLV proviruses are highly polymorphic, but stable DNA elements, that are widespread in the murine genome (24). Under certain circumstances, endogenous MLV can produce infectious virions, a characteristic that differs from HERVs which are generally considered non-infectious (67). Still, also HERVs have been shown to produce viral-like particles in cell culture, cancer and autoimmune diseases (67–69). For example, RNA and protein transcribed from LINE-1 retroelements are packaged into EVs (70). HERV-W Env Syncytin-1 is present on EVs derived from cutaneous T cell lymphoma, BeWo cells and placenta (71, 72).

Accumulating evidence suggests an association of HERV expression with NDs. Some but not all studies demonstrate that HERV-K transcripts are more abundant in ALS patients compared to unaffected individuals (22, 73). Expression appears to be restricted to neurons rather than glia (56). Elevated levels of HERV-K Env peptides in sera and CSF of ALS patients correlated with poor functional performance, suggesting that HERV-K Env contributes to disease progression (74). Impaired ERV repression has also been correlated with Tau pathology (16, 17). Elevated ERV transcripts have been reported for AD (18, 19), PSP (20), behavioral variant frontotemporal dementia (21) and sporadic CJD (75, 76). Brains of bovine spongiform encephalopathy-infected macaques displayed increased ERV transcripts (77). ERV Gag protein and RNA of the retroelement group IAP were also found to co-fractionate with CJD infectivity (78–80). Importantly, HERVs are also upregulated during infection with exogenous pathogens, during inflammation und aging, processes which have been implicated in the progression of NDs (81, 82). While directly being neurotoxic or indirectly activating microglia, our data suggest that ERV de-repression could also contribute to prion-like spreading events.

## Methods

### Molecular Cloning

Murine XPR1 or human XPR1 were amplified from cDNA of N2a NM-GFP^sol^ or HEK NM-GFP^sol^ cells. The open reading frame from of murine XPR1 was tagged with a sequence for the HA-epitope and cloned into PiggyBac expression vector PB510B-1 (System Biosciences) with XbaI and NotI. To generate the phCMV-Syncytin-1-100UTR plasmid, Syncytin-1 cDNA tagged with a V5 epitope sequence (#T0264; GeneCopoeia) was cloned into phCMV-EcoENV (Addgene #15802) using EcoRI and XhoI to replace EcoENV. The 100 bp sequence from 3’-UTR of Syncytin-1 shown to enhance gene expression (83) was amplified using primers (forward: 5’- CCGCTCGAGAGCGGTCGTCGGCCAAC-3’/ reverse: 5’- GAAGATCTCCTTCCCAGCTAGGCTTAGGG-3’) and genomic DNA from MCF-7 cells as template. The sequence was cloned into phCMV-Syncytin-1 using XhoI and BglII restriction sites. phCMV-10A1 (#15805) and pBS-CMV-gagpol (#35614) plasmids were obtained from Addgene.

### Cell lines

N2a cells expressing NM-HA^sol^ or NM-GFP^sol^ and the N2a NM-HA^agg^ clone induced to propagate NM-HA aggregates have been published previously (28, 29). Vero NM-GFP^sol^, Vero Tau-FR^sol^ (-FusionRed), HEK Tau-GFP^sol^ and HEK Tau-FR^sol^ cells have been described previously, as have HEK Tau-GFP^AD^ cells producing Tau-GFP aggregates induced with Alzheimer’s disease brain homogenate (10, 54). All code for human 4R Tau amino acids 243 to 375 containing the mutations P301L and V337M fused to GFP or FR (Evrogen) with an 18-amino acid flexible linker (EFCSRRYRGPGIHRSPTA), as described previously (84). N2a and HEK293T cells were cultured in Opti-MEM (Gibco) supplemented with glutamine, 10 % (v/v) fetal bovine serum (FCS) (PAN-Biotech GmbH) and antibiotics. Vero cells were purchased from CLS (Cell lines service) and cultivated as recommended. Melan-a cells were cultured in RPMI 1640 (Gibco) with 2 mM glutamine, 10 % FCS, P/S and 200 nM PMA. All cells were incubated at 37° C and 5 % CO2. The total numbers of cells and the viability of cells were determined using the Vi-VELL^TM^XR Cell Viability Analyzer (Beckman Coulter). Transfections of cells were performed with either Lipofectamine 2000 or TransIT-2020 / X2 (Mirus) reagents as recommended by the manufacturers.

### Isolation of cortical neurons

Preparation of cortical neurons was performed from the cortices of p13 SWISS pups as described previously (29). Cortical neurons were transduced with lentivirus after 2 d cultivation on 96 well plates or Sarstedt 8 slice chambers. 2 d later, EVs were added to the cortical neurons and incubated for 2 d. The neurons were fixed for microscopy and imaging analysis.

### Production and transduction with lentiviral particles

HEK293T cells were cotransfected with plasmids pRSV-Rev, pMD2.VSV-G, pMDl.g/pRRE, and pRRl.sin.PPT.hCMV.Wpre coding for Tau-FR or NM-GFP. Supernatants were harvested and concentrated with PEG according to published protocols (85). Vero cells and primary neurons were transduced with lentivirus. Stable Vero cell clones with homogenous Tau-GFP/FR expression were produced by limiting dilution cloning (28).

### EV isolation

To prepare EV-depleted medium, fetal calf serum was ultracentrifuged at 100,000 x g, 4° C for 20 h. Medium supplemented with EV-depleted FCS and antibiotics was subsequently filtered using 0.22 and a 0.1 µM filter-sterilization devices (Millipore). For EV isolation, 2-4 x 10^6^ cells were seeded in a T175 flask in 35 ml -depleted medium to be confluent after 3 d. EVs were harvested 3 d post transfection. Cells and cell debris were pelleted by differential centrifugation (300 x g, 10 min; 2,000 x g, 20 min; 16,000 x g, 30 min). The remaining supernatant (conditioned medium) was subjected to ultracentrifugation (UC) (100,000 x g, 1 h) using rotors Ti45 or SW32Ti (Beckman Coulter). The pellet was rinsed with PBS and spun again using rotor SW55Ti (100,000 x g, 1 h). For conditioned medium experiments, cells were transfected with plasmids phCMV-10A1 (Addgene #15805), pBS-CMV-gagpol (Addgene #35614) and phCMV using TransIT. 5 h post transfection, medium was switched to medium with EV-depleted serum. EV-conditioned medium was harvested 3 d post transfection, centrifuged for 10 min at 300 g and subsequently used for aggregate induction assays.

### Aggregate induction assay

Recipient cells were cultured on CellCarrier-96 or 384 black microplate (PerkinElmer) at appropriate cell numbers for 1 h, and then treated with 5-20 µl of prepared samples (isolated EVs, conditioned medium or recombinant NM fibrils). To destroy EVs or vesicles in conditioned medium, samples were sonicated for 5 min at 100 % intensity. For cocultures, recipient and donor cells were seeded at different ratios based on the population doubling time of donor and recipient cells. The total number of cells per well was 3x10^4^. After additional 18 h for NM and 48 h for Tau, cells were fixed with 4 % paraformaldehyde and nuclei were counterstained with Hoechst. Cells were imaged with the automated confocal microscope CellVoyager CV6000 (Yokogawa Inc.) using a 20 x or 40 x objective. Maximum intensity projections were generated from z-stacks. Images from 16 fields per well were taken. At least of 3-4 x 10^3^ cells per well and at least 3 wells per treatment were analyzed.

### Production of recombinant NM

To purify recombinant NM-His, BL21 (DE3) competent *E. coli* were transformed with 100 ng pET vector containing the coding sequence of NM with a C-terminal His-tag under control of the T7 promoter. Five ml of *E. coli* overnight cultures were inoculated into 250 ml LB media containing 100 μg/ ml ampicillin. Cultures were incubated at 37° C, 180 rpm (Multitron, Infors HT), until reaching an OD600 of 0.8. NM-His expression was induced with 1 mM IPTG for 3 h at 37°C, 180 rpm. Centrifuged pellets (10 min, 3000 x g) from 1.5 l bacterial culture were pooled and lysed in 75 ml buffer A (8 M urea, 20 mM imidazole in phosphate buffer) for 1 h at RT. After sonication for 3 x 10 s at 50 % intensity, cell debris was pelleted for 20 min at 10,000 x g and the supernatant was sterile-filtered. NM-His was purified from the supernatant via IMAC using the ÄKTA pure protein purification system (GE Healthcare) together with a 5 ml HisTrap HP His tag protein purification column (GE Healthcare). The supernatant was loaded onto the column initially washed with 25 ml buffer A. After rinsing with 75 ml buffer A, NM-His was eluted using a linear imidazole gradient from 10 mM to 250 mM imidazole (2 % - 50 % buffer B; 8 M urea, 500 mM Imidazole in phosphate buffer). NM-His containing fractions were pooled and concentrated to around 10 % of the initial volume using Vivaspin 20 concentrator columns with a molecular cut-off of 10,000 Da. The protein was desalted using a 5 ml HiTrap Desalting column (GE Healthcare) and sterile-filtered PBS. Protein-containing fractions were pooled and frozen at -80°C.

### Determination of size and number of EVs

ZetaView PMX 110-SZ-488 Nano Particle Tracking Analyzer (Particle Metrix GmbH) was used to determine the size and number of isolated EVs. The instrument captures the movement of extracellular particles by utilizing a laser scattering microscope combined with a video camera. For each measurement, the video data is calculated by the instrument and results in a velocity and size distribution of the particles. For nanoparticle tracking analysis, the Brownian motion of the vesicles from each sample was followed at 22° C with properly adjusted equal shutter and gain. At least six individual measurements of 11 positions within the measurement cell and around 2200 traced particles in each measurement were detected for each sample.

### Sample preparation for mass spectrometry

Cell pellets from 5 replicates, and 6 replicates of EV samples from N2a NM-HA^agg^ subclone s2E at early and late passages were collected for a quantitative proteomics analysis. Cell pellets were lysed in 150 µL SDT buffer (4 % SDS (w/v), 100 mM Tris/HCl pH 7.6, 0.1 M DTT) by homogenization with a dounce tissue grinder and heated 3 min at 95° C. Afterwards, the samples were sonicated 5 times for 30 s with intermediate cooling using a vialtweeter sonifier (amplitude 100 %, duty cycle 50 %; Hielscher, Germany). EV pellets were lysed in 100 µL STET lysis buffer (150 mM NaCl, 50 mM TrisHCl pH 7.5, 2 mM EDTA, 1 % Triton X-100) on ice for 30 min with intermediate vortexing. Cell debris and undissolved material was removed by centrifugation at 16,000 × g for 5 min. The protein concentrations were measured using the colorimetric 660 nm assay (Thermo Fisher Scientific, US). For cell lysates, the assay solution was supplemented with the ionic detergent compatibility reagent (Thermo Fisher Scientific, US). A protein amount of 30 µg per sample for cell lysates and 10 µg for EV lysates was subjected to proteolytic digestion using the filter aided sample preparation (FASP) protocol (86) with 30 kD Vivacon spin filters (Sartorius, Germany). Proteolytic peptides were desalted by stop and go extraction (STAGE) with C18 tips (87). The purified peptides were dried by vacuum centrifugation. Digestions of cell lysates and EVs were dissolved in 40 or 20 µl of 0.1 % formic acid, respectively.

### LC-MS/MS analysis

Samples were analyzed by LC-MS/MS for relative label free protein quantification. A peptide amount of approximately 1 µg per sample was separated on a nanoLC system (EASY-nLC 1000, Proxeon – part of Thermo Scientific, US) using in-house packed C18 columns (50 cm or 30 cm x 75 µm ID, ReproSil-Pur 120 C18-AQ, 1.9 µm, Dr. Maisch GmbH, Germany) with a binary gradient of water (A) and acetonitrile (B) containing 0.1% formic acid at 50° C column temperature and a flow rate of 250 nl/ min. Cell lysates were separated on a 50 cm column using a gradient of 250 min length, whereas a 183 min gradient on a 30 cm column was used for EV samples (250 min. gradient: 0 min., 2 % B; 5 min., 5 % B; 185 min., 25 % B; 230 min., 35 % B; 250 min., 60 % B; 183 min. gradient: 0 min., 2 % B; 3:30 min., 5 % B; 137:30 min., 25 % B; 168:30 min., 35 % B; 182:30 min., 60 % B). The nanoLC was coupled online via a nanospray flex ion source (Proxeon – part of Thermo Scientific, US) equipped with a PRSO-V2 column oven (Sonation, Germany) to a Q-Exactive mass spectrometer (Thermo Scientific, US). Full MS spectra were acquired at a resolution of 70,000. The top 10 peptide ions were chosen for Higher-energy C-trap Dissociation (HCD) with a normalized collision energy of 25 %. Fragment ion spectra were acquired at a resolution of 17,500. A dynamic exclusion of 120 s was used for peptide fragmentation.

### Data analysis and label free quantification

The raw data was analyzed by the software Maxquant (maxquant.org, Max-Planck Institute Munich) version and 1.5.5.1 (88). The MS data was searched against a FASTA database of *Mus musculus* from UniProt including also non-reviewed entries supplemented with databases of lentiviruses and murine leukemia viruses (download: December 09^th^ 2017, 52041 + 712 + 43 entries). Trypsin was defined as protease. Two missed cleavages were allowed for the database search. The option first search was used to recalibrate the peptide masses within a window of 20 ppm. For the main search peptide and peptide fragment mass tolerances were set to 4.5 and 20 ppm, respectively. Carbamidomethylation of cysteine was defined as static modification. Acetylation of the protein N-term as well as oxidation of methionine were set as variable modifications. The false discovery rate for both peptides and proteins was adjusted to less than 1 %. Label free quantification (LFQ) of proteins required at least two ratio counts of razor peptides. Only unique and razor peptides were used for quantification. The LFQ values were log2^ transformed and a two-sided Student’s t-test was used to evaluate statistically significant changed abundance of proteins between cell lysates from passages 16 and 7 as well as EV lysates from passages 15 and 6. A p-value less than 5 % was set as significance threshold. Additionally, a permutation based false discovery rate estimation was used to account for multiple hypotheses (89).

### OptiPrep density gradient

For separating EVs and viruses, the discontinuous iodixanol gradient in 1.2 % increments ranging from 6 to 18% were prepared as previously described (40). The 100,000 x g pellet from 1050 ml culture supernatant (30 x T175 flasks) was resuspended in 1 ml PBS and overlaid onto the gradient. The gradient was subjected to high-speed centrifugation at 100,000 x g for 2 h at 4° C using a SW41Ti rotor (Beckman Coulter). 12 fractions of 1ml each were collected from the top of the gradient, diluted with PBS in 5 ml, and centrifuged at 100,000 x g for 1 h at 4° C. The pelleted fractions were resuspended in 100 µl PBS, and then used for further experiments. The reverse transcriptase activity of the viruses was measured by using the colorimetric Reverse Transcriptase Assay (Roche).

### Electron Microscopy (EM)

EM imaging of EV and virus preparations were performed as previously described (42). Briefly, the 100,000 x g pellets were fixed in 2 % paraformaldehyde, loaded on glow discharged Formvar / carbon-coated EM grids (Plano GmbH), contrasted in uranyl-oxalat (pH 7) for 5 min and embedded in uranyl-methylcellulose for 5 min. Samples were examined using a JEOL JEM-2200FS transmission electron microscope at 200 kV (JEOL).

### Infectivity assay

The infectivity assay was performed as previously described (57). Briefly, melan-a cells were exposed to conditioned medium from different cell clones at either low or high passages in the presence of 4 µg Polybrene/ ml for 24 h. The medium was then replaced with normal culture medium. After 6 d, cells were lysed for Western blot analysis for the existence of retroviral Env and Gag proteins.

### Drug Treatment

The treatment of cells with Amprenavir (10 µM; Selleckchem), Lopinavir (10 µM; Selleckchem) and DMSO was performed for 72 h in EV-depleted medium in T175 flasks. Afterwards, the total numbers of viable cells and the viability upon drug treatments were determined using the Vi-VELL^TM^XR Cell Viability Analyzer (Beckman Coulter). EVs were isolated from the conditioned medium via ultracentrifugation and processed for the aggregate induction assay as described above. Coculture and EV aggregate induction for NM were performed in the absence of the drugs, whereas all cell-based assays were performed in the presence of drugs at the same concentration as the pretreatment. To inhibit methyltransferase, N2a NM-HA^agg^ (EP) donor cells were treated for 3 d with methyltransferase inhibitors 5-Azacytidine (Aza) (Sigma-Aldrich), Decitabine (Dec) (Sigma-Aldrich) or DMSO as solvent control. Subsequently, the cells were cultured in the absence of the drugs for 5 d. Afterwards, the treated donor cells were cocultured with recipient cells as described above for their aggregate inducing capacity and analyzed by Western blot for MLV Env and Gag expression levels. To increase DNA methylation, the N2aq NM-HA^agg^ (LP) donor clone were treated with methyl group donors L-methionine (L-M), Betaine (B), Choline chloride (CC) or medium control for 6 d. MLV Env and Gag protein levels were analyzed by Western blot. Subsequently, cells were cocultured with recipient cells for 16 h. The percentage of aggregate containing recipient cells was compared to the percentage of aggregate bearing recipients cocultured with solvent-treated donors.

### Neutralization assay

To block MLV Env on the surface of the donor cell clone and on EVs, mAb83A25, which reacts with almost all members of MLVs (90) was incubated with either EVs or donor cells in serial dilutions for 1 h at 37°C with rotation at 20 rpm. Afterwards, the donor cells were mixed with recipient cells and EVs were added to the pre-seeded recipient cells for 1 d.

### Transfection with siRNAs or plasmids

To transiently knock-down the upregulated specific MLV Env and Gag genes in N2a NM- HA^agg^ donors, custom-designed Silencer select siRNAs (Thermo Fisher) against AAO37244.2 (*env*) and AID54952 (*gag*) were used. Pre-designed siRNAs against murine XPR1 and mCat-1 were used to knock-down both genes. For transfection, cells were pre-seeded on 6 well plate 1 d before at 2x10^5^ cells/well. The next day, cells were transfected with a final concentration of 60 nM siRNA or control siRNA using lipofectamine RNAiMax diluted in Opti-MEM for 30 min before addition to cells. After 2-3 d, cells were collected for aggregate induction assays, qRT-PCR or Western blot analysis.

### qRT-PCR

Validation of changes of different genes in cell pellets were carried out by real-time quantitative polymerase chain reaction (qPCR). First, total RNAs from cell pellets were isolated using the RNeasy Mini Kit or RNeasy Lipid Tissue Mini Kit (Qiagen). RNA concentration and quality were determined with Agilent 2100 Bioanalyzer System. RNAs were reversely transcribed to cDNA using the iScript^TM^ cDNA Synthesis Kit (Bio-Rad). To determine mRNA expression levels of murine *env* (AAO37244.2) and *gag* (AID54952), custom designed TaqMan assays and for murine *pan-env*, *xpr1*, *mcat-1* and *gapdh* as housekeeping control pre-designed TagMan assays were used. For detection of the MLV subclasses, mRNA expression levels were measured using SYBR Green assays (Applied Biosystems). For quantitative real-time PCR, the fold change was calculated with the ΔΔCT method. Following primers were used: *Ecotropic* MLV (forward:5’- AGGCTGTTCCAGAGATTGTG -3’; reverse: 5’-CCGGGGCAGACATAGAATCC-3’), *Xenotropic* MLV (forward:5’-GGAGCCTACCAAGCACTCA-3’; reverse: 5’- GGCAGAGGTATGGTTGGAGTA-3’), *GAPDH* (forward:5’- GCACAGTCAAGGCCGAGAAT-3’; reverse: 5’-GCCTTCTCCATGGTGGTGAA -3’).

### Western blotting

For Western blot analysis, protein concentrations were measured by Quick Start^TM^ Bradford Protein assay (Bio-Rad) using the plate reader Fluostar Omega BMG (BMG Labtech) and the corresponding MARS Data Analysis Software (BMG Labtech). Proteins were separated on NuPAGE®Novex® 4-12 % Bis-Tris Protein Gels (Life Technologies) followed by transfer onto a PVDF membrane (GE Healthcare) in a wet blotting chamber. Western blot analysis was performed using rat hybridoma anti-MLV Env mAb83A25 (90) (1:10; kindly provided by L.H. Evans, Rocky Mountain Laboratories, MT); goat anti- xenotropic MLV virus antibody ABIN457298 (1:1000, antibodies-online); mouse anti-MLV gag ab100970 (1:1000, Abcam); rat anti-HA 3F10 (1:1000, Roche); mouse anti-GAPDH 6C5 (1:5000, Abcam); mouse anti-Actin C4 (1:5000; MP Biomedical); rabbit anti-Tau ab64193 (1:1000, Abcam); mouse anti PiT-2 B-4 (1:1000; Santa Cruz Biotechnology); rabbit anti-V5 D3H8Q (1:1000, Cell signaling); mouse anti-Alix (1:1000, BD Bioscience); mouse anti-Hsc/Hsp70 N27F3-4 (1:1000, ENZO); rabbit anti-Flotillin1 ab 133497 (1:1000, Abcam). The membrane was incubated with Pierce^TM^ ECL Western blotting substrate (Thermo Fisher Scientific) according to the manufactureŕs recommendations and imaged with the Imaging system Fusion FX (Vilber Lourmat).

### Automated image Analysis

The image analysis was performed using the CellVoyager Analysis support software (CV7000 Analysis Software; Version 3.5.1.18). An image analysis routine was developed for single cell segmentation and aggregate identification (Yokogawa Inc.) The total number of cells was determined based on the Hoechst signal, and recipient cells were detected by their GFP/FR signal. Respective green or red aggregates were identified via morphology and intensity characteristics. The percentage of recipient cells with aggregated NM-GFP or Tau-FR was calculated as the number of aggregate-positive cells per total recipient cells set to 100 %.

### Immunofluorescence staining and confocal microscopy analysis

Cells were fixed in 4 % paraformaldehyde. For HA-staining cells were rinsed with PBS, permeabilized in 0.1 % Triton X-100, blocked in 2 % goat serum in PBS and incubated for with rabbit anti-HA ab9110 (Abcam) antibody diluted 1:500 in blocking solution for 2 h at RT. After three washing steps with PBS, cells were incubated for 1 h with Alexa Fluor 647- conjugated secondary antibody, while nuclei were counterstained with 4 µg/ml Hoechst 33342 (Molecular Probes). 384 well plates were scanned with CellVoyager CV6000 (Yokogawa Inc.). Confocal laser scanning microscopy was performed on a Zeiss LSM 800 laser-scanning microscope with Airyscan (Carl Zeiss) and analyzed via Zen2010 (ZenBlue, Zeiss).

### Statistical analysis

All analyses were performed using the Prism 6.0 (GraphPad Software v.7.0c). Statistical inter-group comparisons were performed using the one-way ANOVA with a Bonferroni post-test or Student’s unpaired t-test. The confidence interval in both tests was 95 %, p values smaller than 0.1 were considered significant. All experiments were performed in at least triplicates (EV experiments) or at least sextuplicates (coculture experiments) and repeated at least two times independently with similar results. Measurements were taken from distinct samples. At least 6000 cells were analyzed for quantitative analysis. Shown are the mean and the error bar representing the standard deviation (SD).

## Acknowledgements

We thank Leonard Henry “Pug” Evans for generously sharing anti-MLV antibodies. We are grateful to Paolo Salomoni and Dan Ehninger for critical reading of the manuscript. The light microcopy (LMF) and laboratory automation facilities (LAT) of the DZNE Bonn were used for image acquisition.

## Funding information

This work was funded by the Helmholtz Portfolio “Wirkstoffforschung”, the “Deutsche Forschungsgemeinschaft” (DFG, German Research Foundation) under Germany’s Excellence Strategy within the framework of the “Munich Cluster for Systems Neurology” (EXC 2145 SyNergy– ID 390857198), by the German Ministry for Education and research through grants CLINSPECT-M (FKZ161L0214C) and JPND PMG-AD (01ED2002B).

The funders had no role in study design, data collection and analysis, decision to publish, or preparation of the manuscript.

## Financial competing interests

S. Liu, S.A. Müller, S.F. Lichtenthaler, P. Denner and I.M. Vorberg hold pending patent applications for “HERV inhibitors for use in treating tauopathies”: “US Patent Application No. 17/640,119 based on PCT International application No. PCT/EP2020/074809, claiming priority to “European Application No. 19195304.1”.

## Supplementary figures

**Suppl. Figure 1. Increased aggregate induction by donors that have been in culture for prolonged time. a**. N2a NM cell culture model. N2a cells expressing soluble NM-HA (NM-HA^sol^, upper left image) or soluble NM-GFP (NM-GFP^sol^, upper right) (28). N2a NM-HA^sol^ cells were exposed to recombinant NM fibrils and donor clone N2a NM-HA^agg^ s2E was established that consistently produces NM-HA aggregates (NM-HA^agg^, lower images). For simplicity, we call the clone N2a NM-HA^agg^ (30). Shown are early (P7) and late passages of the clone (P16). **b.** Cocultures of N2a NM-GFP^sol^ with N2a NM-HA^agg^ cells of early and late passage. As controls, recipients were also cocultured with N2a NM- HA^sol^ cells. **c.** Cryopreserved N2a NM-HA^agg^ cells of late passage (P21) retain NM aggregate high inducing activity. Donor cells were frozen at passage 1. Donors of different passaging history were subsequently defrosted and cultured for less than 6 passages (thereafter termed “EP” for “early passage” and “LP” for “late passage”, respectively). Donors were cocultured with N2a NM-GFP^sol^ cells and recipients with NM-GFP^agg^ were detected 16 h later. As control, recipients were cultured with N2a NM-HA^sol^ cells. **d.** Confocal images of recipient cells exposed to conditioned medium from donors of early or late passage. To destroy vesicles, medium was sonicated. **e.** Quantitative analysis of aggregate induction by conditioned medium shown in **e**. **f.** Confocal images of recipient cells exposed to isolated EVs from donors of early or late passage. **g.** Quantitative analysis of aggregate induction by EVs. To destroy vesicles, EVs were sonicated. Insets show higher magnifications of cells. All data are shown as the means ± SD from three (g) or six (c, e) replicate cell cultures. Three (c, e, g) independent experiments were carried out with similar results. P-values calculated by two-tailed unpaired Student’s t-test (e, g) or one-way ANOVA (c). Source data are provided as a Source Data file.

**Suppl. Figure 2. Upregulation of endogenous retrovirus is not caused by aggregate induction. a.** N2a NM-HA^sol^ cells were exposed to 1 µM recombinant NM fibrils (monomer equivalent) and cells were subsequently cultured for 15 days. Untreated cells served as controls. NM-HA was detected using anti-HA antibody. **b.** Quantitative analysis of recipient cells with induced aggregates. NM-HA aggregate induction was monitored 72 h post exposure. **c.** Expression of eco- and xeno-/polytropic MLV was detected by RT-PCR. **d.** N2a NM-GFP^sol^ cells were exposed to 1 µM recombinant NM fibrils (monomer equivalent) and cells were subsequently cultured for 15 days. Untreated cells served as controls. **e.** Quantitative analysis of recipient cells with induced aggregates. NM-GFP aggregate induction was monitored 72 h post exposure. **f.** Expression of eco- and xeno-/polytropic MLV was detected by RT-PCR. **g.** Recipient N2a NM-GFP^sol^ cells were exposed to conditioned medium (CM) from early or late passage donors in the presence or absence of Vectofusin-1. Aggregate induction was assessed 16 h later. **h.** Quantitative analysis of g. All data are shown as the means ± SD from three (c, f, h) or six (b, e) replicate cell cultures. Three (b-c, e-f, h) independent experiments were carried out with similar results. P-values calculated by two-tailed unpaired Student’s t-test. ns: non-significant. Source data are provided as a Source Data file.

**Suppl. Figure 3. HIV Integrase, reverse transcriptase and HCV protease inhibitors fail to inhibit intercellular aggregate induction. a.** Effect of HIV protease inhibitors on preexisting NM-HA aggregates in donor cells. Shown are the percentages of donor cells with aggregates upon exposure to different concentrations of drugs. **b.** Experimental scheme. N2a NM-HA^agg^ and N2a NM-GFP^sol^ cells were cocultured in the presence of HIV or HCV inhibitors. Effect of different concentrations of inhibitors on the percentage of donor and recipient cells harboring NM-HA^agg^ or NM-GFP^agg^, respectively. % cells with aggregates were normalized to control cocultures only treated with DMSO. All data are shown as the means ± SD from three (a-b) replicate cell cultures. Three (a-b) independent experiments were carried out with similar results. Source data are provided as a Source Data file.

**Suppl. Figure 4. MLV receptor knock-down has no influence on aggregate induction by recombinant NM fibrils. a.** Recipient N2a NM-GFP^sol^ cells were transfected with two siRNAs against the XPR1 receptor or one siRNA against mCat-1. As controls, recipients were transfected with non-silencing siRNA. Two days later, recipients were exposed to µm recombinant NM fibril (monomer equivalent). Recipients with NM-GFP^agg^ were quantified 16 h post exposure. **b.** Recipient HEK NM-GFP^sol^ cells were transfected with a plasmid coding for the N2a XPR1 receptor or empty plasmid for 2 d. Afterwards, pretreated recipients were exposed to 10 µm recombinant NM fibril (monomer equivalent). As controls, recipients N2a NM-GFP^sol^ were induced with NM fibril in the same way. Recipients with NM-GFP^agg^ were quantified 16 h post exposure. All data are shown as the means ± SD from three (a) or six (b) replicate cell cultures. Three (a-b) independent experiments were carried out with similar results. P-values calculated by two-tailed unpaired Student’s t-test (b) or one-way ANOVA (a). ns: non-significant. Source data are provided as a Source Data file.

## Notes

### Competing Interest Statement

S. Liu, S.A. Mueller, S.F. Lichtenthaler, P. Denner and I. M. Vorberg hold pending patent applications for HERV inhibitors for use in treating tauopathies: US Patent Application No. 17/640,119 based on PCT International application No. PCT/EP2020/074809, claiming priority to European Application No. 19195304.1.

## References

1. La Joie R, Visani AV, Baker SL, Brown JA, Bourakova V, Cha J, Chaudhary K, Edwards L, Iaccarino L, Janabi M, Lesman-Segev OH, Miller ZA, Perry DC, O’Neil JP, Pham J, Rojas JC, Rosen HJ, Seeley WW, Tsai RM, Miller BL, Jagust WJ, Rabinovici GD. 2020. Prospective longitudinal atrophy in Alzheimer’s disease correlates with the intensity and topography of baseline tau-PET. Sci Transl Med 12.

2. Valori CF, Neumann M. 2021. Contribution of RNA/DNA Binding Protein Dysfunction in Oligodendrocytes in the Pathogenesis of the Amyotrophic Lateral Sclerosis/Frontotemporal Lobar Degeneration Spectrum Diseases. Front Neurosci 15:724891.

3. Ghetti B, Oblak AL, Boeve BF, Johnson KA, Dickerson BC, Goedert M. 2015. Invited review: Frontotemporal dementia caused by microtubule-associated protein tau gene (MAPT) mutations: a chameleon for neuropathology and neuroimaging. Neuropathol Appl Neurobiol 41:24–46.

4. Vogel JW, Iturria-Medina Y, Strandberg OT, Smith R, Levitis E, Evans AC, Hansson O, Alzheimer’s Disease Neuroimaging I, Swedish BioFinder S. 2020. Spread of pathological tau proteins through communicating neurons in human Alzheimer’s disease. Nat Commun 11:2612.

5. Braak H, Braak E. 1991. Neuropathological stageing of Alzheimer-related changes. Acta Neuropathol 82:239–59.

6. Goedert M, Clavaguera F, Tolnay M. 2010. The propagation of prion-like protein inclusions in neurodegenerative diseases. Trends Neurosci 33:317–25.

7. Clavaguera F, Bolmont T, Crowther RA, Abramowski D, Frank S, Probst A, Fraser G, Stalder AK, Beibel M, Staufenbiel M, Jucker M, Goedert M, Tolnay M. 2009. Transmission and spreading of tauopathy in transgenic mouse brain. Nat Cell Biol 11:909–13.

8. Kaufman SK, Sanders DW, Thomas TL, Ruchinskas AJ, Vaquer-Alicea J, Sharma AM, Miller TM, Diamond MI. 2016. Tau Prion Strains Dictate Patterns of Cell Pathology, Progression Rate, and Regional Vulnerability In Vivo. Neuron 92:796–812.

9. Brunello CA, Merezhko M, Uronen RL, Huttunen HJ. 2020. Mechanisms of secretion and spreading of pathological tau protein. Cell Mol Life Sci 77:1721–1744.

10. Liu S, Hossinger A, Heumuller SE, Hornberger A, Buravlova O, Konstantoulea K, Muller SA, Paulsen L, Rousseau F, Schymkowitz J, Lichtenthaler SF, Neumann M, Denner P, Vorberg IM. 2021. Highly efficient intercellular spreading of protein misfolding mediated by viral ligand-receptor interactions. Nat Commun 12:5739.

11. Wells JN, Feschotte C. 2020. A Field Guide to Eukaryotic Transposable Elements. Annu Rev Genet 54:539–561.

12. Belshaw R, Watson J, Katzourakis A, Howe A, Woolven-Allen J, Burt A, Tristem M. 2007. Rate of recombinational deletion among human endogenous retroviruses. J Virol 81:9437–42.

13. Geis FK, Goff SP. 2020. Silencing and Transcriptional Regulation of Endogenous Retroviruses: An Overview. Viruses 12.

14. Kury P, Nath A, Creange A, Dolei A, Marche P, Gold J, Giovannoni G, Hartung HP, Perron H. 2018. Human Endogenous Retroviruses in Neurological Diseases. Trends Mol Med 24:379–394.

15. Kassiotis G. 2014. Endogenous retroviruses and the development of cancer. J Immunol 192:1343–9.

16. Guo C, Jeong HH, Hsieh YC, Klein HU, Bennett DA, De Jager PL, Liu Z, Shulman JM. 2018. Tau Activates Transposable Elements in Alzheimer’s Disease. Cell Rep 23:2874–2880.

17. Protasova MS, Gusev FE, Grigorenko AP, Kuznetsova IL, Rogaev EI, Andreeva TV. 2017. Quantitative Analysis of L1-Retrotransposons in Alzheimer’s Disease and Aging. Biochemistry (Mosc) 82:962–971.

18. Frost B, Hemberg M, Lewis J, Feany MB. 2014. Tau promotes neurodegeneration through global chromatin relaxation. Nat Neurosci 17:357–66.

19. Dembny P, Newman AG, Singh M, Hinz M, Szczepek M, Kruger C, Adalbert R, Dzaye O, Trimbuch T, Wallach T, Kleinau G, Derkow K, Richard BC, Schipke C, Scheidereit C, Stachelscheid H, Golenbock D, Peters O, Coleman M, Heppner FL, Scheerer P, Tarabykin V, Ruprecht K, Izsvak Z, Mayer J, Lehnardt S. 2020. Human endogenous retrovirus HERV-K(HML-2) RNA causes neurodegeneration through Toll-like receptors. JCI Insight 5.

20. Sun W, Samimi H, Gamez M, Zare H, Frost B. 2018. Pathogenic tau-induced piRNA depletion promotes neuronal death through transposable element dysregulation in neurodegenerative tauopathies. Nat Neurosci 21:1038–1048.

21. Phan K, He Y, Fu Y, Dzamko N, Bhatia S, Gold J, Rowe D, Ke YD, Ittner LM, Hodges JR, Piguet O, Kiernan MC, Halliday GM, Kim WS. 2021. Pathological manifestation of human endogenous retrovirus K in frontotemporal dementia. Commun Med (Lond) 1.

22. Douville R, Liu J, Rothstein J, Nath A. 2011. Identification of active loci of a human endogenous retrovirus in neurons of patients with amyotrophic lateral sclerosis. Ann Neurol 69:141–51.

23. Bhardwaj N, Montesion M, Roy F, Coffin JM. 2015. Differential expression of HERV-K (HML-2) proviruses in cells and virions of the teratocarcinoma cell line Tera-1. Viruses 7:939–68.

24. Kozak CA. 2014. Origins of the endogenous and infectious laboratory mouse gammaretroviruses. Viruses 7:1–26.

25. Young GR, Eksmond U, Salcedo R, Alexopoulou L, Stoye JP, Kassiotis G. 2012. Resurrection of endogenous retroviruses in antibody-deficient mice. Nature 491:774–8.

26. Yu P, Lubben W, Slomka H, Gebler J, Konert M, Cai C, Neubrandt L, Prazeres da Costa O, Paul S, Dehnert S, Dohne K, Thanisch M, Storsberg S, Wiegand L, Kaufmann A, Nain M, Quintanilla-Martinez L, Bettio S, Schnierle B, Kolesnikova L, Becker S, Schnare M, Bauer S. 2012. Nucleic acid-sensing Toll-like receptors are essential for the control of endogenous retrovirus viremia and ERV-induced tumors. Immunity 37:867–79.

27. King OD, Gitler AD, Shorter J. 2012. The tip of the iceberg: RNA-binding proteins with prion-like domains in neurodegenerative disease. Brain Res 1462:61–80.

28. Krammer C, Kryndushkin D, Suhre MH, Kremmer E, Hofmann A, Pfeifer A, Scheibel T, Wickner RB, Schatzl HM, Vorberg I. 2009. The yeast Sup35NM domain propagates as a prion in mammalian cells. Proc Natl Acad Sci U S A 106:462–7.

29. Hofmann JP, Denner P, Nussbaum-Krammer C, Kuhn PH, Suhre MH, Scheibel T, Lichtenthaler SF, Schatzl HM, Bano D, Vorberg IM. 2013. Cell-to-cell propagation of infectious cytosolic protein aggregates. Proc Natl Acad Sci U S A 110:5951–6.

30. Liu S, Hossinger A, Hofmann JP, Denner P, Vorberg IM. 2016. Horizontal Transmission of Cytosolic Sup35 Prions by Extracellular Vesicles. MBio 7.

31. Nissen KK, Laska MJ, Hansen B, Terkelsen T, Villesen P, Bahrami S, Petersen T, Pedersen FS, Nexo BA. 2013. Endogenous retroviruses and multiple sclerosis-new pieces to the puzzle. BMC Neurol 13:111.

32. Lieber MM, Todaro GJ. 1973. Spontaneous and induced production of endogenous type-C RNA virus from a clonal line of spontaneously transformed BALB-3T3. Int J Cancer 11:616–27.

33. Alais S, Simoes S, Baas D, Lehmann S, Raposo G, Darlix JL, Leblanc P. 2008. Mouse neuroblastoma cells release prion infectivity associated with exosomal vesicles. Biol Cell 100:603–15.

34. Aaronson SA, Dunn CY. 1974. Endogenous C-type viruses of BALB-c cells: frequencies of spontaneous and chemical induction. J Virol 13:181–5.

35. Rasheed S, Bruszewski J, Rongey R, Roy-Burman P, Charman HP, Gardner MB. 1976. Spontaneous release of endogenous ecotropic type C virus from rat embryo cultures. J Virol 18:799–803.

36. Kim JW, Closs EI, Albritton LM, Cunningham JM. 1991. Transport of cationic amino acids by the mouse ecotropic retrovirus receptor. Nature 352:725–8.

37. Kozak CA. 2010. The mouse "xenotropic" gammaretroviruses and their XPR1 receptor. Retrovirology 7:101.

38. Li M, Xu F, Muller J, Hearing VJ, Gorelik E. 1998. Ecotropic C-type retrovirus of B16 melanoma and malignant transformation of normal melanocytes. Int J Cancer 76:430–6.

39. Radek C, Bernadin O, Drechsel K, Cordes N, Pfeifer R, Strasser P, Mormin M, Gutierrez-Guerrero A, Cosset FL, Kaiser AD, Schaser T, Galy A, Verhoeyen E, Johnston ICD. 2019. Vectofusin-1 Improves Transduction of Primary Human Cells with Diverse Retroviral and Lentiviral Pseudotypes, Enabling Robust, Automated Closed-System Manufacturing. Hum Gene Ther 30:1477–1493.

40. Dettenhofer M, Yu XF. 1999. Highly purified human immunodeficiency virus type 1 reveals a virtual absence of Vif in virions. J Virol 73:1460–7.

41. Ribet D, Harper F, Esnault C, Pierron G, Heidmann T. 2008. The GLN family of murine endogenous retroviruses contains an element competent for infectious viral particle formation. J Virol 82:4413–9.

42. Thery C, Amigorena S, Raposo G, Clayton A. 2006. Isolation and characterization of exosomes from cell culture supernatants and biological fluids. Curr Protoc Cell Biol Chapter 3:Unit 3 22.

43. Risser R, Horowitz JM, McCubrey J. 1983. Endogenous mouse leukemia viruses. Annu Rev Genet 17:85–121.

44. Rowe HM, Trono D. 2011. Dynamic control of endogenous retroviruses during development. Virology 411:273–87.

45. Strick R, Strissel PL, Baylin SB, Chiappinelli KB. 2016. Unraveling the molecular pathways of DNA-methylation inhibitors: human endogenous retroviruses induce the innate immune response in tumors. Oncoimmunology 5:e1122160.

46. Chiappinelli KB, Strissel PL, Desrichard A, Li H, Henke C, Akman B, Hein A, Rote NS, Cope LM, Snyder A, Makarov V, Budhu S, Slamon DJ, Wolchok JD, Pardoll DM, Beckmann MW, Zahnow CA, Merghoub T, Chan TA, Baylin SB, Strick R. 2015. Inhibiting DNA Methylation Causes an Interferon Response in Cancer via dsRNA Including Endogenous Retroviruses. Cell 162:974–86.

47. Ramos MP, Wijetunga NA, McLellan AS, Suzuki M, Greally JM. 2015. DNA demethylation by 5-aza-2’-deoxycytidine is imprinted, targeted to euchromatin, and has limited transcriptional consequences. Epigenetics Chromatin 8:11.

48. Schultz A, Rein A. 1985. Maturation of murine leukemia virus env proteins in the absence of other viral proteins. Virology 145:335–9.

49. Rein A, Mirro J, Haynes JG, Ernst SM, Nagashima K. 1994. Function of the cytoplasmic domain of a retroviral transmembrane protein: p15E-p2E cleavage activates the membrane fusion capability of the murine leukemia virus Env protein. J Virol 68:1773–81.

50. Oshima M, Muriaux D, Mirro J, Nagashima K, Dryden K, Yeager M, Rein A. 2004. Effects of blocking individual maturation cleavages in murine leukemia virus gag. J Virol 78:1411–20.

51. Feher A, Boross P, Sperka T, Miklossy G, Kadas J, Bagossi P, Oroszlan S, Weber IT, Tozser J. 2006. Characterization of the murine leukemia virus protease and its comparison with the human immunodeficiency virus type 1 protease. J Gen Virol 87:1321–30.

52. Battini JL, Rasko JE, Miller AD. 1999. A human cell-surface receptor for xenotropic and polytropic murine leukemia viruses: possible role in G protein-coupled signal transduction. Proc Natl Acad Sci U S A 96:1385–90.

53. Bamunusinghe D, Liu Q, Lu X, Oler A, Kozak CA. 2013. Endogenous gammaretrovirus acquisition in Mus musculus subspecies carrying functional variants of the XPR1 virus receptor. J Virol 87:9845–55.

54. Chakraborty P, Riviere G, Liu S, de Opakua AI, Dervisoglu R, Hebestreit A, Andreas LB, Vorberg IM, Zweckstetter M. 2021. Co-factor-free aggregation of tau into seeding-competent RNA-sequestering amyloid fibrils. Nat Commun 12:4231.

55. Antony JM, van Marle G, Opii W, Butterfield DA, Mallet F, Yong VW, Wallace JL, Deacon RM, Warren K, Power C. 2004. Human endogenous retrovirus glycoprotein-mediated induction of redox reactants causes oligodendrocyte death and demyelination. Nat Neurosci 7:1088–95.

56. Li W, Lee MH, Henderson L, Tyagi R, Bachani M, Steiner J, Campanac E, Hoffman DA, von Geldern G, Johnson K, Maric D, Morris HD, Lentz M, Pak K, Mammen A, Ostrow L, Rothstein J, Nath A. 2015. Human endogenous retrovirus-K contributes to motor neuron disease. Sci Transl Med 7:307ra153.

57. Pothlichet J, Heidmann T, Mangeney M. 2006. A recombinant endogenous retrovirus amplified in a mouse neuroblastoma is involved in tumor growth in vivo. Int J Cancer 119:815–22.

58. Ottina E, Levy P, Eksmond U, Merkenschlager J, Young GR, Roels J, Stoye JP, Tuting T, Calado DP, Kassiotis G. 2018. Restoration of Endogenous Retrovirus Infectivity Impacts Mouse Cancer Models. Cancer Immunol Res 6:1292–1300.

59. Gregory DA, Lyddon TD, Johnson MC. 2013. Multiple Gag domains contribute to selective recruitment of murine leukemia virus (MLV) Env to MLV virions. J Virol 87:1518–27.

60. D’Souza V, Summers MF. 2005. How retroviruses select their genomes. Nat Rev Microbiol 3:643–55.

61. Iguchi Y, Eid L, Parent M, Soucy G, Bareil C, Riku Y, Kawai K, Takagi S, Yoshida M, Katsuno M, Sobue G, Julien JP. 2016. Exosome secretion is a key pathway for clearance of pathological TDP-43. Brain 139:3187–3201.

62. Sproviero D, La Salvia S, Giannini M, Crippa V, Gagliardi S, Bernuzzi S, Diamanti L, Ceroni M, Pansarasa O, Poletti A, Cereda C. 2018. Pathological Proteins Are Transported by Extracellular Vesicles of Sporadic Amyotrophic Lateral Sclerosis Patients. Front Neurosci 12:487.

63. Leblanc P, Alais S, Porto-Carreiro I, Lehmann S, Grassi J, Raposo G, Darlix JL. 2006. Retrovirus infection strongly enhances scrapie infectivity release in cell culture. EMBO J 25:2674–85.

64. Leblanc P, Hasenkrug K, Ward A, Myers L, Messer RJ, Alais S, Timmes A, Priola SA. 2012. Co-infection with the friend retrovirus and mouse scrapie does not alter prion disease pathogenesis in susceptible mice. PLoS One 7:e30872.

65. Krasemann S, Neumann M, Luepke JP, Grashorn J, Wurr S, Stocking C, Glatzel M. 2012. Persistent retroviral infection with MoMuLV influences neuropathological signature and phenotype of prion disease. Acta Neuropathol 124:111–26.

66. Maksakova IA, Romanish MT, Gagnier L, Dunn CA, van de Lagemaat LN, Mager DL. 2006. Retroviral elements and their hosts: insertional mutagenesis in the mouse germ line. PLoS Genet 2:e2.

67. Lower R, Lower J, Frank H, Harzmann R, Kurth R. 1984. Human teratocarcinomas cultured in vitro produce unique retrovirus-like viruses. J Gen Virol 65 (Pt 5):887–98.

68. Ono M, Kawakami M, Ushikubo H. 1987. Stimulation of expression of the human endogenous retrovirus genome by female steroid hormones in human breast cancer cell line T47D. J Virol 61:2059–62.

69. Christensen T, Dissing Sorensen P, Riemann H, Hansen HJ, Moller-Larsen A. 1998. Expression of sequence variants of endogenous retrovirus RGH in particle form in multiple sclerosis. Lancet 352:1033.

70. Bowers EC, Motta A, Knox K, McKay BS, Ramos KS. 2022. LINE-1 Cargo and Reverse Transcriptase Activity Profiles in Extracellular Vesicles from Lung Cancer Cells and Human Plasma. Int J Mol Sci 23.

71. Laukkanen K, Saarinen M, Mallet F, Aatonen M, Hau A, Ranki A. 2020. Cutaneous T-Cell Lymphoma (CTCL) Cell Line-Derived Extracellular Vesicles Contain HERV-W-Encoded Fusogenic Syncytin-1. J Invest Dermatol 140:1466–1469 e4.

72. Vargas A, Zhou S, Ethier-Chiasson M, Flipo D, Lafond J, Gilbert C, Barbeau B. 2014. Syncytin proteins incorporated in placenta exosomes are important for cell uptake and show variation in abundance in serum exosomes from patients with preeclampsia. FASEB J 28:3703–19.

73. Tam OH, Ostrow LW, Gale Hammell M. 2019. Diseases of the nERVous system: retrotransposon activity in neurodegenerative disease. Mob DNA 10:32.

74. Arru G, Mameli G, Deiana GA, Rassu AL, Piredda R, Sechi E, Caggiu E, Bo M, Nako E, Urso D, Mariotto S, Ferrari S, Zanusso G, Monaco S, Sechi G, Sechi LA. 2018. Humoral immunity response to human endogenous retroviruses K/W differentiates between amyotrophic lateral sclerosis and other neurological diseases. Eur J Neurol 25:1076–e84.

75. Muotri AR, Chu VT, Marchetto MC, Deng W, Moran JV, Gage FH. 2005. Somatic mosaicism in neuronal precursor cells mediated by L1 retrotransposition. Nature 435:903–10.

76. Jeong BH, Lee YJ, Carp RI, Kim YS. 2010. The prevalence of human endogenous retroviruses in cerebrospinal fluids from patients with sporadic Creutzfeldt-Jakob disease. J Clin Virol 47:136–42.

77. Greenwood AD, Vincendeau M, Schmadicke AC, Montag J, Seifarth W, Motzkus D. 2011. Bovine spongiform encephalopathy infection alters endogenous retrovirus expression in distinct brain regions of cynomolgus macaques (Macaca fascicularis). Mol Neurodegener 6:44.

78. Murdoch GH, Sklaviadis T, Manuelidis EE, Manuelidis L. 1990. Potential retroviral RNAs in Creutzfeldt-Jakob disease. J Virol 64:1477–86.

79. Akowitz A, Sklaviadis T, Manuelidis L. 1994. Endogenous viral complexes with long RNA cosediment with the agent of Creutzfeldt-Jakob disease. Nucleic Acids Res 22:1101–7.

80. Manuelidis L, Sklaviadis T, Akowitz A, Fritch W. 1995. Viral particles are required for infection in neurodegenerative Creutzfeldt-Jakob disease. Proc Natl Acad Sci U S A 92:5124–8.

81. Licastro F, Porcellini E. 2021. Activation of Endogenous Retrovirus, Brain Infections and Environmental Insults in Neurodegeneration and Alzheimer’s Disease. Int J Mol Sci 22.

82. Romer C. 2021. Viruses and Endogenous Retroviruses as Roots for Neuroinflammation and Neurodegenerative Diseases. Front Neurosci 15:648629.

83. Kitao K, Tanikaga T, Miyazawa T. 2019. Identification of a post-transcriptional regulatory element in the human endogenous retroviral syncytin-1. J Gen Virol 100:662–668.

84. Woerman AL, Aoyagi A, Patel S, Kazmi SA, Lobach I, Grinberg LT, McKee AC, Seeley WW, Olson SH, Prusiner SB. 2016. Tau prions from Alzheimer’s disease and chronic traumatic encephalopathy patients propagate in cultured cells. Proc Natl Acad Sci U S A 113:E8187–E8196.

85. Follenzi A, Naldini L. 2002. HIV-based vectors. Preparation and use. Methods Mol Med 69:259–74.

86. Wisniewski JR, Zougman A, Nagaraj N, Mann M. 2009. Universal sample preparation method for proteome analysis. Nat Methods 6:359–62.

87. Rappsilber J, Ishihama Y, Mann M. 2003. Stop and go extraction tips for matrix-assisted laser desorption/ionization, nanoelectrospray, and LC/MS sample pretreatment in proteomics. Anal Chem 75:663–70.

88. Cox J, Hein MY, Luber CA, Paron I, Nagaraj N, Mann M. 2014. Accurate proteome-wide label-free quantification by delayed normalization and maximal peptide ratio extraction, termed MaxLFQ. Mol Cell Proteomics 13:2513–26.

89. Tusher VG, Tibshirani R, Chu G. 2001. Significance analysis of microarrays applied to the ionizing radiation response. Proc Natl Acad Sci U S A 98:5116–21.

90. Evans LH, Morrison RP, Malik FG, Portis J, Britt WJ. 1990. A neutralizable epitope common to the envelope glycoproteins of ecotropic, polytropic, xenotropic, and amphotropic murine leukemia viruses. J Virol 64:6176–83.

