## Supplementary material for "Endogenous retroviruses promote prion-like spreading of proteopathic seeds": Suppl. Fig

Author contributions: S.L., S.-E.H., S.F.L. and I.V. designed research; S.L., S.-E.H., A. H., O.B., S.A.M. performed research; M. N. contributed human samples; S.F.L., S.A.M.P.D. and I.V. analyzed data; S. L., S.-E.H. and I. V. wrote the paper.

Suppl. Figure 1

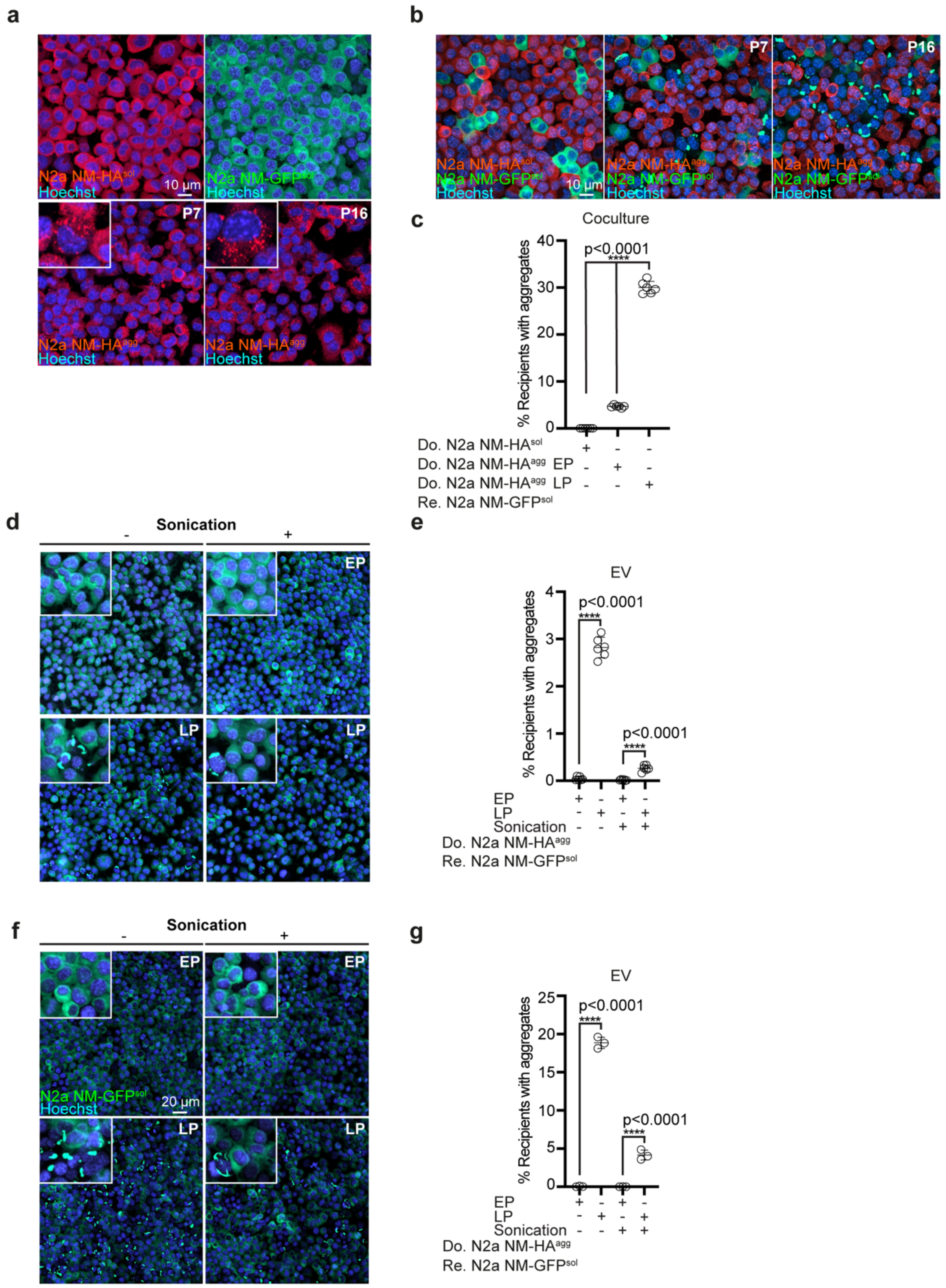

Suppl. Figure 2

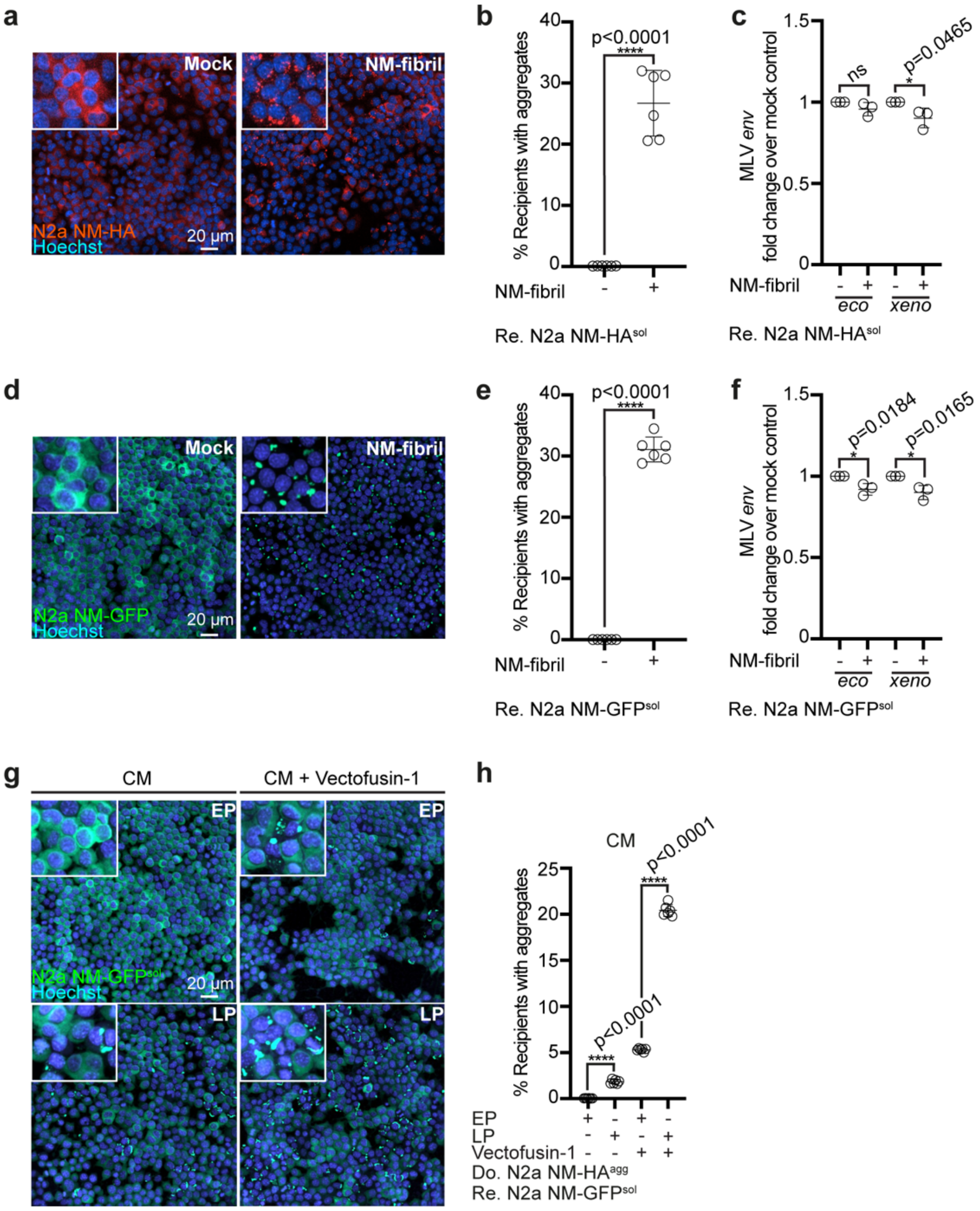

Suppl. Figure 3

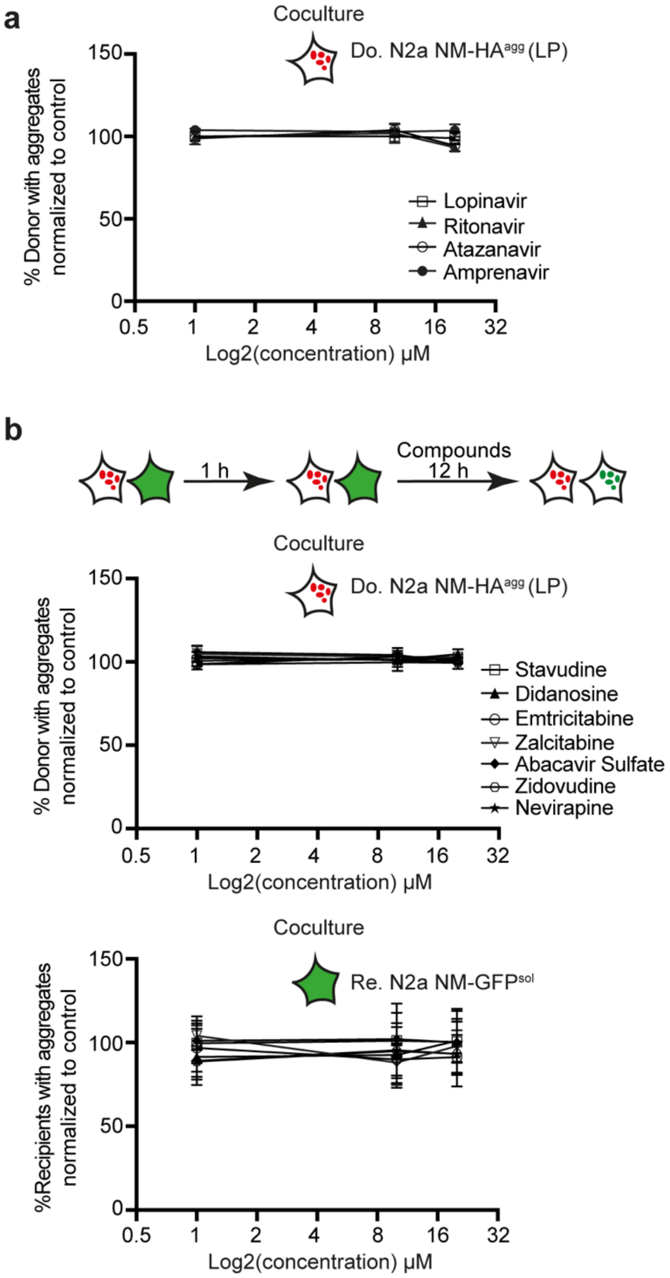

Suppl. Figure 4

**a**

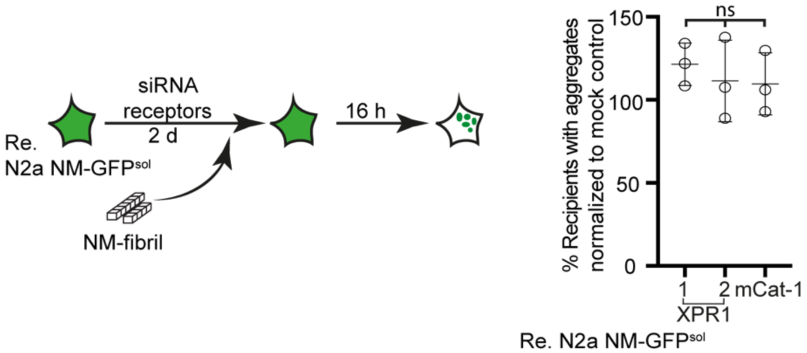


**b**

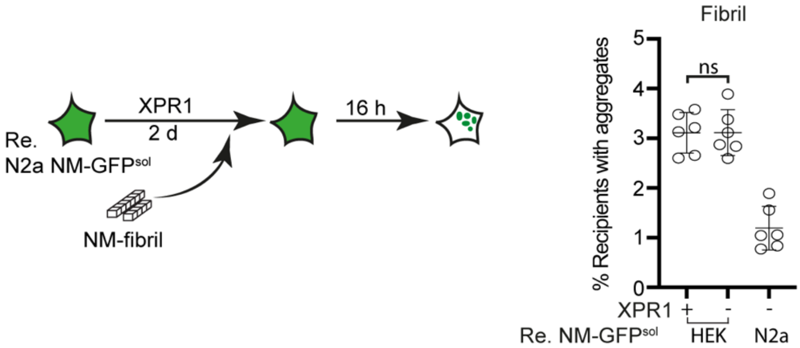
